# Meta-analysis and experimental re-evaluation of the Boyle van ‘t Hoff relation with osmoregulation modelled by linear elastic principles and ion-osmolyte leakage

**DOI:** 10.1101/2022.03.05.483010

**Authors:** Dominic J Olver, Iqra Azam, James D Benson

## Abstract

In this study we challenge the paradigm of using the Boyle van ’t Hoff (BvH) relation to relate cell size as a linear function of inverse extracellular osmotic pressure for short time periods (~5 to 30 mins). We present alternative models that account for mechanical resistance (turgor model) and ion-osmolyte leakage (leak model), which is not accounted for by the BvH relation. To test the BvH relation and the alternative models, we conducted a meta-analysis of published BvH datasets, as well as new experiments using a HepG2 cell line. Our meta-analysis showed that the BvH relation may be assumed of the hypertonic region but cannot be assumed *a priori* over the hyper- and hypotonic region. Both alternative models perform better than the BvH relation but are nearly indistinguishable when plotted. The return to isotonic conditions plot indicated neither alternative model accurate predicts return volumes for HepG2 cells. However, a combined turgor-leak model accurately predicts both the BvH plot and the return to isotonic conditions plot. Moreover, this turgor-leak model provides a facile method to estimate the membrane-cortex Young’s modulus and the cell membrane permeability to intracellular ions/osmolytes during periods of osmotic challenge, and predicts a novel passive method of volume regulation without the need for ion pumps.

## 1 Introduction

Cell shape and size are governed by an interplay of hydrostatics, cytoskeletal dynamics, membrane contractility, membrane and cortical tension, and electrical and chemical potential [1–4]. Controlling cell shape and size is critical for cell growth [5,6], development [7], locomotion [8], and function [9]. As water is nearly incompressible, intracellular water content is directly related to cell shape and size, but also integral to molecular crowding [10], mass flux behaviour [11], and cellular mechanics [8,12]. All eukaryotic membranes, with few exceptions, are highly permeable to water which, depending on the cell, constitutes 10 to 90% of cell volume [1]. Since water is several orders of magnitude more permeable in the lipid membrane than ions [13–15], equilibrium water volume is governed by differences in intracellular osmotic compounds, extracellular tonicity, and osmosis [16,17]. Much recent attention has been paid to mechanobiological models that rely on accurate estimates of intracellular water content. These estimates improve our understanding of long and short term volume regulation [18,19], electroosmotics [20], cell-size regulation [6], cellular migration [21], cellular aging [22], and cancer modelling [23,24].

The Boyle van ’t Hoff (BvH) relation (or model) has been used widely to determine the intracellular water volume (osmotically active volume) and the non-osmotic volume of the cell for more than 100 years [25–28]. The BvH relation models cell volume as proportional to inverse osmotic pressure, and in this context, it is a linear model. However, there are alternative nonlinear models for cells that do not appear to follow the BvH relation [1,29,30]. Alternative models include osmoregulation (regulation of intracellular osmotic conditions and by extension, volumetric response) through either mechanical resistance [31], regulating internal solute concentration [18], or accounting for the chemical potential of proteins in the cell at high concentrations [32]. Presence of osmoregulation may lead to inaccurate estimates of osmotically active volume and thus inaccurate models. On the other hand, identification of mechanisms for osmoregulation improves estimates for osmotically active volume along with selection of appropriate model for volumetric responses to osmotic dynamics [4,11,15,18].

### 1.1 Boyle van ‘t Hoff Relation

The BvH relation arose from an analogous form of Boyle’s gas law applied to cells in ideal dilute solutions [27,28]. Osmotic pressure *Π*, and intracellular solution volume 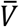, are related through the van ’t Hoff equation 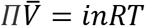, where *n* is the number of mols of solute, *i* is the van ’t Hoff factor for the respective solute, *R* is the gas constant, and *T* is temperature [27,28]. At equilibrium, the intracellular osmotic pressure *Π*_in_, will be equal to extracellular osmotic pressure *Π*_ext_, and thus, for a given external osmotic pressure, there is a corresponding solution volume such that 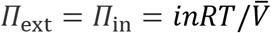. Osmolarity, *c*, is typically used to calculate osmotic pressure. However, a theoretical inconsistency arises at the limit when 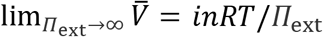, then 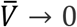: since 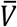 is a solution volume and therefore includes the nonzero volume of ions and osmolytes inside the cell, there is a contradiction since 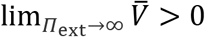. Using solvent volume *W* corrects for this inconsistency and thus linearly relating osmotic pressure to osmolality, *π*, and intracellular solvent volume (commonly referred to as osmotically active volume) is theoretically preferable [33,34]. To change between solvent volume and solvent mass, the density of water, *ρ*, may be used to modify the van ’t Hoff equation to *ΠW* = *inRT*/*ρ*.

The cell volume, *V*, can be defined in terms of osmotically active volume (*W*) and non-osmotic (also referred to as non-solvent, non-osmotically active or osmotically inactive) volume, *V*_b_, which includes solutes such as salts and proteins, and non-soluble volume or “solids” of the cell, as well as the bound water forming hydration shells around these items [27,33], such that *V* = *W* + *V*_b_. Note that *V*_b_ does not typically include lipid membrane permeable solutes experimentally introduced into the environment such as urea or dimethyl sulfoxide. The BvH relation assumes that there is no hydrostatic resistance at the membrane, the lipid membrane is semi-permeable, and *inRT*/*ρ* is constant. Thus,

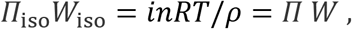

where the subscript iso indicates a value at isotonic conditions [27,28]. The isotonic reference point is not strictly necessary since any osmotic condition that is ideal and dilute can be used for the above relation, but the isotonic point is useful for identifying osmotically active volume and non-osmotic volume for the cell in isotonic conditions (i.e. its normal rest state). Rearranging and combining, *Π*_iso_*W*_iso_ = *Π W*, with the total cell volume model, we obtain

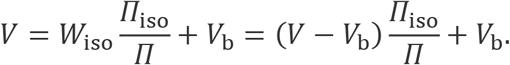

Normalizing with respect to isotonic volume *V*_iso_ provides a non-dimensionalized relation in the form, *ν* = *w* + *b*, where *ν* = *V*/*V*_iso_ is the relative cell volume, *w* = *W*/*V*_iso_ is the osmotically active fraction, and *b* = *k*_b_/*V*_iso_ is the non-osmotic fraction of the cell. Nondimensionalizing the BvH relation enables direct comparison between cells regardless of cell type, size, and condition [11]. Nondimensionalizing osmotic pressure with isotonic osmotic pressure, *Π*_iso_, results in the same ratio as relative inverse osmolality, since *Π*_iso_/*Π* = *RTπ*_iso_/*RTπ* = *π*_iso_/*π*, for osmolality *π*. The non-dimensional BvH relation takes the form:

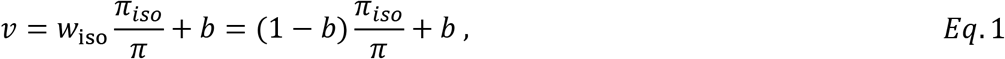

for which *w*_iso_ and *b* are typically linearly regressed with in a BvH plot when isotonic volume is not explicitly known before treatment, and (1 – *b*) is regressed when the isotonic volume is explicitly known [11,25,26]. To be clear, volume is linearly related to *inverse* osmolality (or *inverse* osmotic pressure). The fitted parameters, *w*_iso_, and *b*, inform important models including mass transfer models that enable predictions of volumetric change over time as a function of extracellular osmotic conditions [11,35,36]. Furthermore, despite differences in fitted parameters, osmotic pressure can be estimated as a linear function of either osmolality or osmolarity, thus *Eq*. 1 retains its linear form irrespective of using *Π*_iso_/*Π*, *π*_iso_/*π*, or *c*_iso_/*c* [27,28,34].

The BvH relation describes the relationship between equilibrated cell water volume and osmotic pressure of the environmental solution. Both the extracellular and intracellular solutions are assumed to be ideal and dilute. To determine the water (osmotically active) volume and solids (osmotically inactive) volume of the cell, one can plot a linear regression of *V*/*V*_iso_ with respect to *Π*_iso_/*Π* with the fitted intercept being the non-osmotic fraction of the cell, *b*, while the slope of the regression may be interpreted as the osmotically active fraction of the cell *w*_iso_. This plot is commonly referred to as a Boyle van ’t Hoff plot or Ponder’s plot (Figure 1; [37–39]).

**Figure 1.**
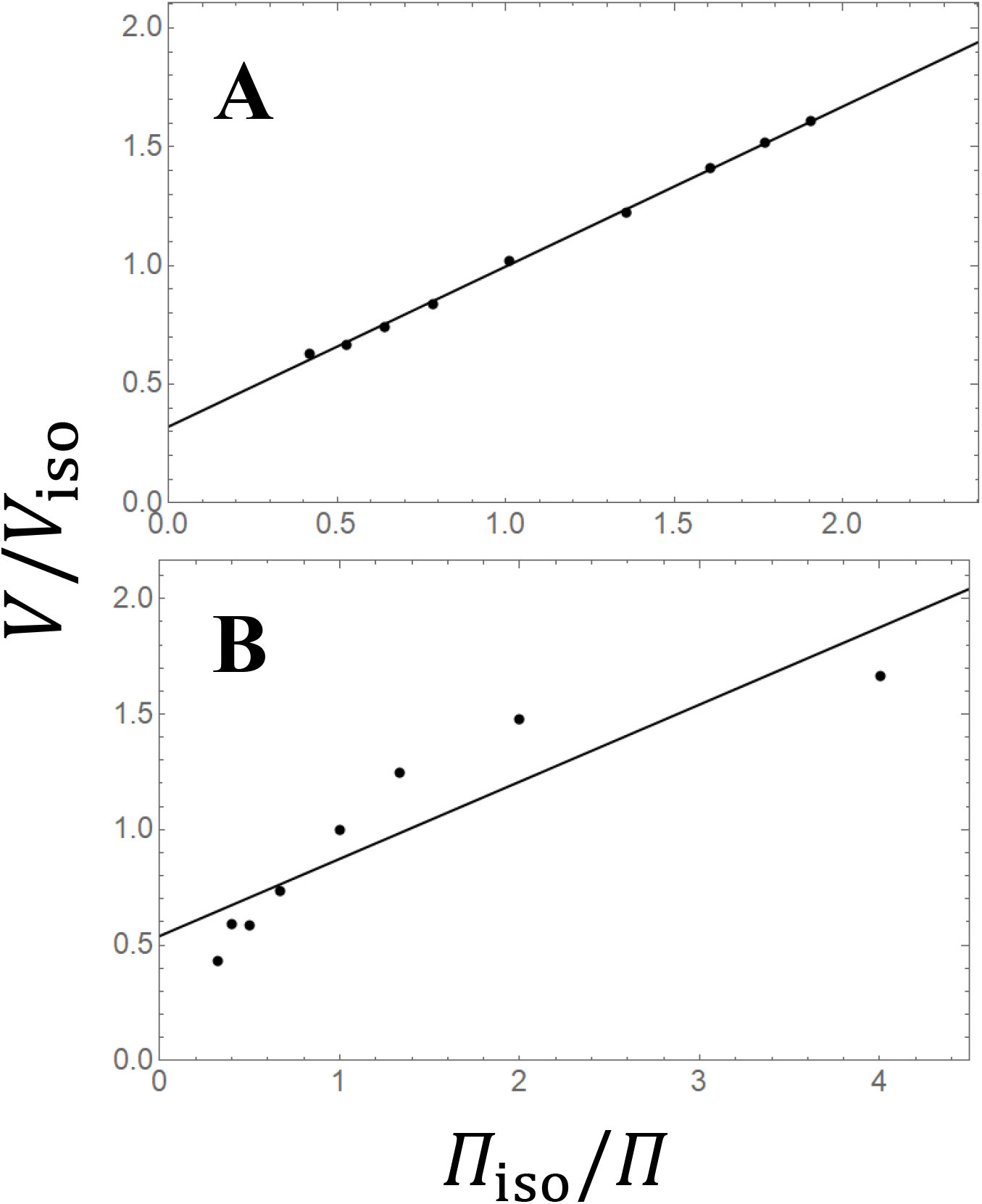
Two Boyle van ’t Hoff plots are presented above with the BvH relation plotted as the black line. Plot **A** is an example of an ideal osmometer with data extracted from Finan et al. (2009) [38]. Plot **B** is an example of a non-ideal osmometer with data extracted from Freeman et al. (1966) [108]. Note the difference of inverse relative osmotic pressure range between the two datasets.

The Boyle van ’t Hoff relation makes two additional assumptions concerning the cell [27,28,32]. First, that the cell membrane is impermeable to ions and osmolytes, and second, that the cell membrane has no significant resistive mechanical force. Cells that satisfy these conditions and appear to behave linearly in the BvH plot are so-called “ideal osmometers” (or “perfect osmometers”; Figure 1 A). The BvH relation also assumes an ideal dilute solution in the cell itself. Prickett and colleagues point out non-ideal non-dilute solutions stand as a particular issue for erythrocytes in hypertonic conditions, and thus the BvH relation is not valid for estimating non-osmotic volume for erythrocytes [32] in very hypertonic conditions. However, non-ideal non-dilute solutions for cells in general may pose a serious issue for the BvH relation validity. Furthermore, organisms have evolved methods of osmoregulation such as the Amoeba [40], notochordal cells [41], astrocytes [42], and many more [1]. Methods of osmoregulation are not taken into account in the BvH relation. Indeed, there are multiple examples of cells that do not appear to behave as ideal osmometers (Figure 1 B; [1,29,30]).

### 1.2 HepG2 Cells

The liver is an organ that carries out a broad range of important functions such as detoxification, metabolism, and homeostasis. Hepatocytes, in particular, are the major parenchymal cells in liver and are employed as an important model for a variety of research questions in the field of pharmacological and toxicological research [43]. Human hepatoma cell line HepG2 are widely used as a biological model of hepatocytes and are of particular interest [44–47]. Despite their wide use, to our knowledge there currently does not exist a Boyle van ’t Hoff plot of HepG2 cells and little is known concerning the underlying mechanisms of volume regulation [48]. Indeed, HepG2 cells are mechanosensitive to osmotic/mechanical perturbations, and are known to change permeability of ions as a function of osmotic and mechanical perturbations [48–51]. Such behaviours make HepG2 cells difficult to study in the traditional context of the BvH relation since underlying assumptions of the BvH relation are not necessarily met, even for short time frames. Development of a systematic method to identifying underlying mechanisms of osmoregulation would not just be useful for obtaining important cellular parameters and understanding underlying cellular mechanics, but also enable improved cell physiological modelling of HepG2 cells. Such a development would be of particular importance for cell types that are not well behaved in the context of the BvH relation such as golden hamster oocytes [52], MDCK cells [42], and Hela cells [53].

In this study, we conduct a meta-analysis and test the general validity of the Boyle van ’t Hoff relation and find it is not an appropriate general model relating relative volume to relative inverse osmotic pressure across hyper- and hypotonic conditions. However, the Boyle van ’t Hoff relation is not found to be inappropriate when fit to just the hypertonic region. We present two generalized nonlinear models, the leak model, and the turgor model, modelling for passive osmolyte regulation and volumetric mechanical resistance respectively. Informed by the meta-analysis, we experimentally identify the mechanism of osmoregulation for HepG2 cells, a human hepatocellular carcinoma cell line. The combined the turgor-leak model, accurately predicts both BvH plot and return to isotonic conditions plot indicating HepG2 cells osmoregulate by a combination of membrane-cortex mechanical resistance and ion-osmolyte leakage. The turgor-leak model uniquely predicts a novel mechanism of time dependent volume regulation by use of mechanical resistance to passively change ion concentration over time, without the need for active pumps.

## 2 Results

### 2.1 Meta-Anaylsis

A linear regression, representing the BvH relation (*Eq*. 1), was performed for each of the 28 respective datasets of Group 1 (nondimensionalized hyper- and hypertonic datasets with 6 or more mean data points). The Durbin Watson (DW) score was calculated for each dataset with 6 or more mean data points and the null hypothesis of no autocorrelation (α=0.05) was tested. Of the 28 datasets, 5 datasets (17.9%) had an associated DW score lower than the respective critical value. An exact binomial test was performed (*n*=28,α=0.05) with a significant result (*p*=0.0117) indicating the number of observed rejected null hypotheses (17.9%) was significantly greater than expected (5%). A downward bend is apparent when plotting the residuals of each of the linear regressions together (Figure 2). More information of each particular dataset is provided in Supplementary Material (SM1).

**Figure 2.**
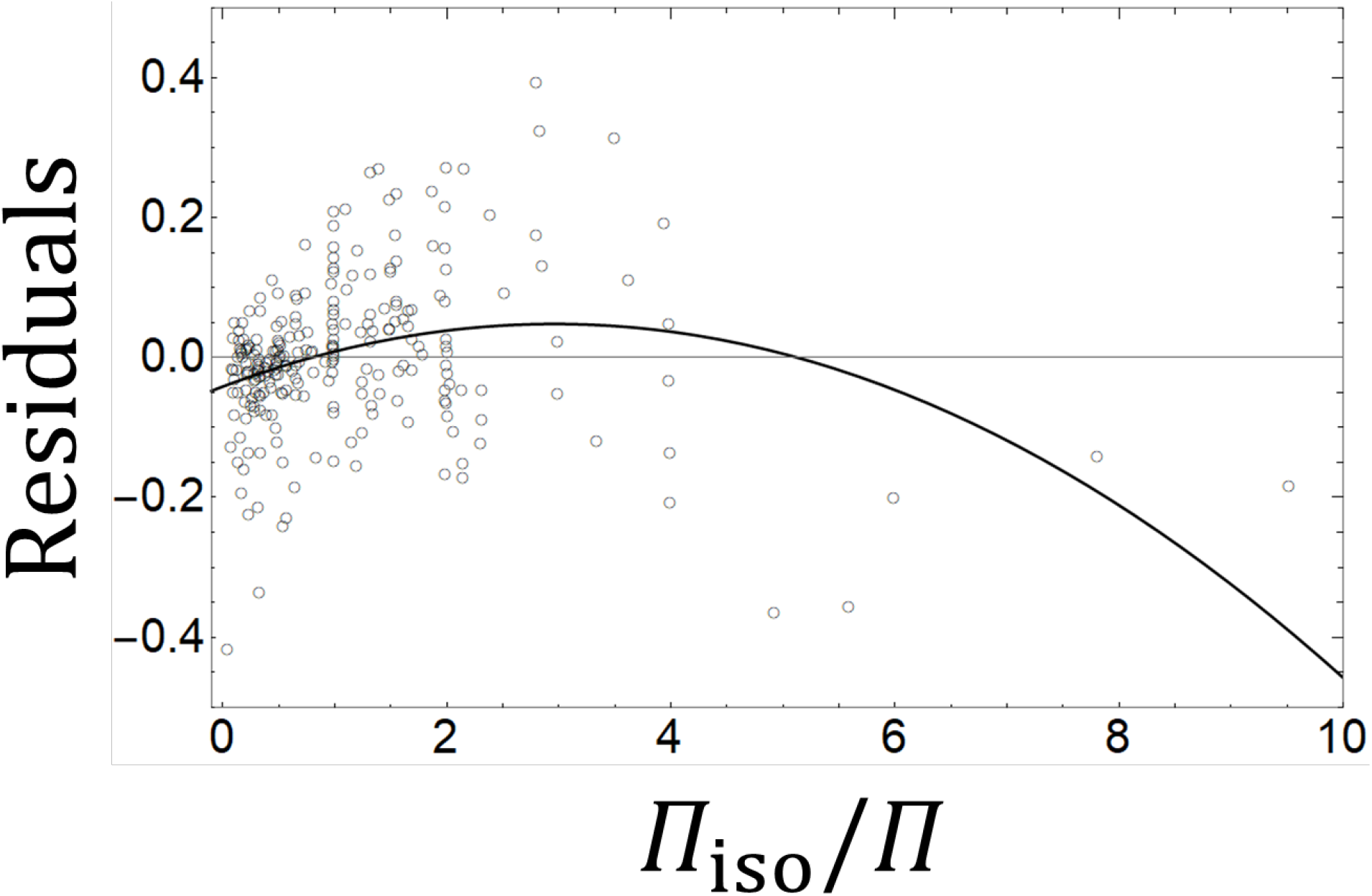
Fitted residuals of the BvH relation for Group 1 including 28 of datasets and 247 data points in across both the hyper- and hypotonic regions. Datasets used were required to have 6 or more data points and ranged from at least 0.5 to 2.0 relative inverse osmotic pressure. Thick black line is a fitted second order polynomial providing a visual aid. Data is originally in terms of inverse osmotic pressure, osmolarity or osmolality. See SM1 for more information on each dataset.

A linear regression was performed for each of the 22 respective datasets of Group 2 (nondimensionalized hypertonic datasets with 6 or more mean data points). The DW score was calculated for each dataset with 6 or more mean data points in the hypertonic region and the null hypothesis of no autocorrelation (α=0.05) was tested. Of the 22 datasets, 2 datasets (9.1%) had an associated DW score lower than their respective critical value. An exact binomial test was performed (*n*=22, α=0.05) with a non-significant result (*p*=0.302) indicating the number of observed rejected null hypotheses (9.1%) was not significantly greater than expected (5%). When plotting the residuals of each linear regression together, there is no visual trend apparent (Figure 3). More information of each particular dataset from Group 2 is provided in Supplementary Material (SM2).

**Figure 3.**
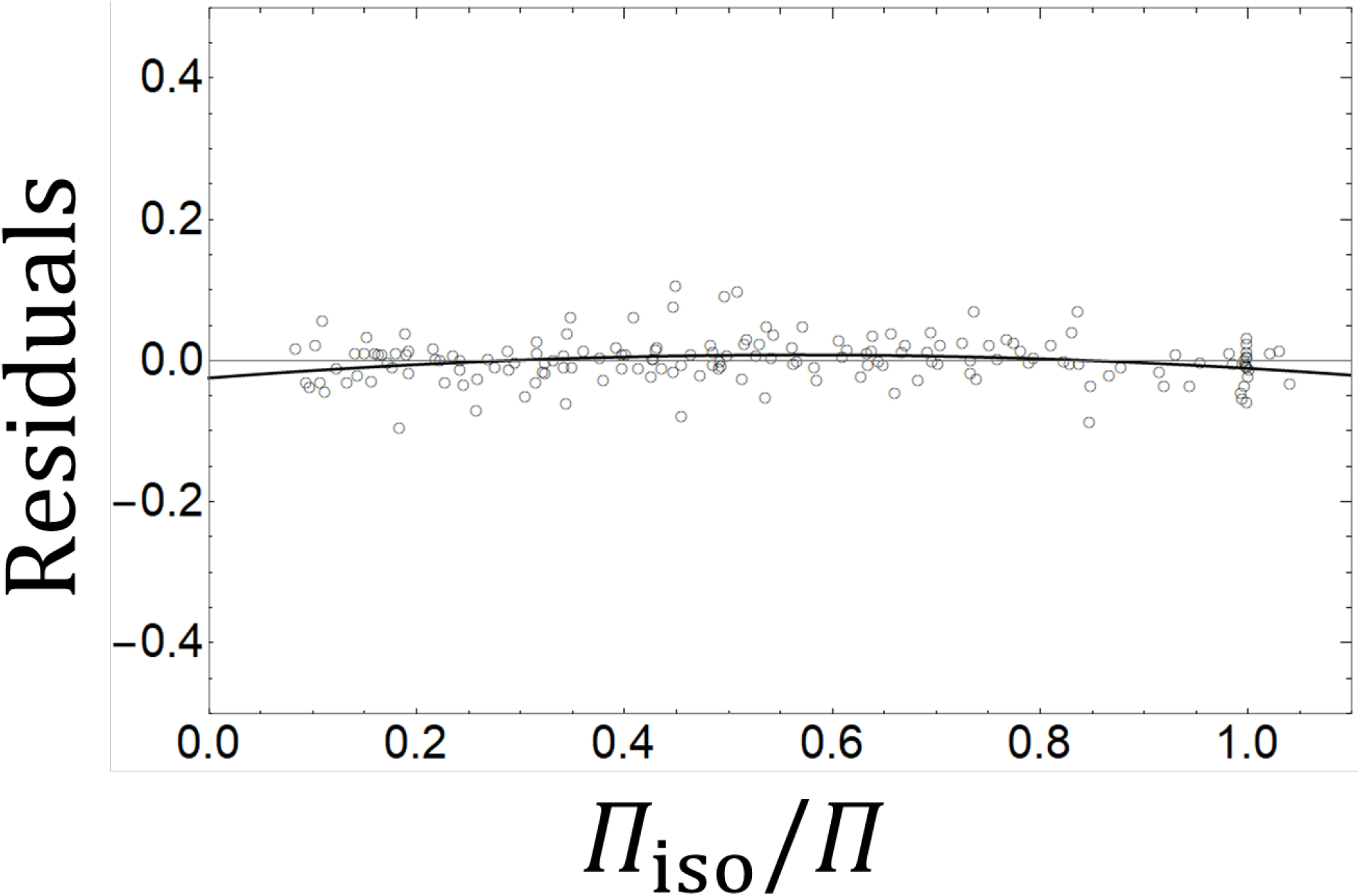
Group 2 BvH relation (hypertonic) Fitted residuals of the BvH relation for 22 of datasets and 167 data points in the hypertonic region. Thick black line is a fitted second order polynomial providing a visual aid. See SM2 for more information on each dataset.

To determine if the fitted constants, *w*_iso_ and *b*, change when regressing the full domain or just the hypertonic region of the BvH plot, a linear regression was performed on 44 datasets in Group 3 (nondimensionalized hyper- and hypotonic datasets) with and without the occlusion of the hypotonic region. The fitted parameters are shown in Figure 4. For regressions including both the hyper- and hypotonic regions, the median *w*_iso_ is 0.516 with an interquartile range (IQR) of 0.316 (range between 25% quartile to 75% quartile) while fitting to just the hypertonic region the median *w*_iso_ is 0.698 (IQR=0.286) and is significantly different (Mann-Whiteny U=626, *p*=0.0043). Again, for the regression over the full domain, the median *b* is 0.471 (IQR=0.302) while fitting to just the hypertonic region the median *b* is 0.303 (IQR=0.271) and is significantly different (Mann-Whiteny U=1266, *p*=0.0130). For Group 3, the BvH relation had a downward bend in the residual plot when using both the hypertonic and hypotonic data (Figure 5) but did not have a noticeable downward bend in the residual plot when using just the hypertonic data (Figure 6).

**Figure 4.**
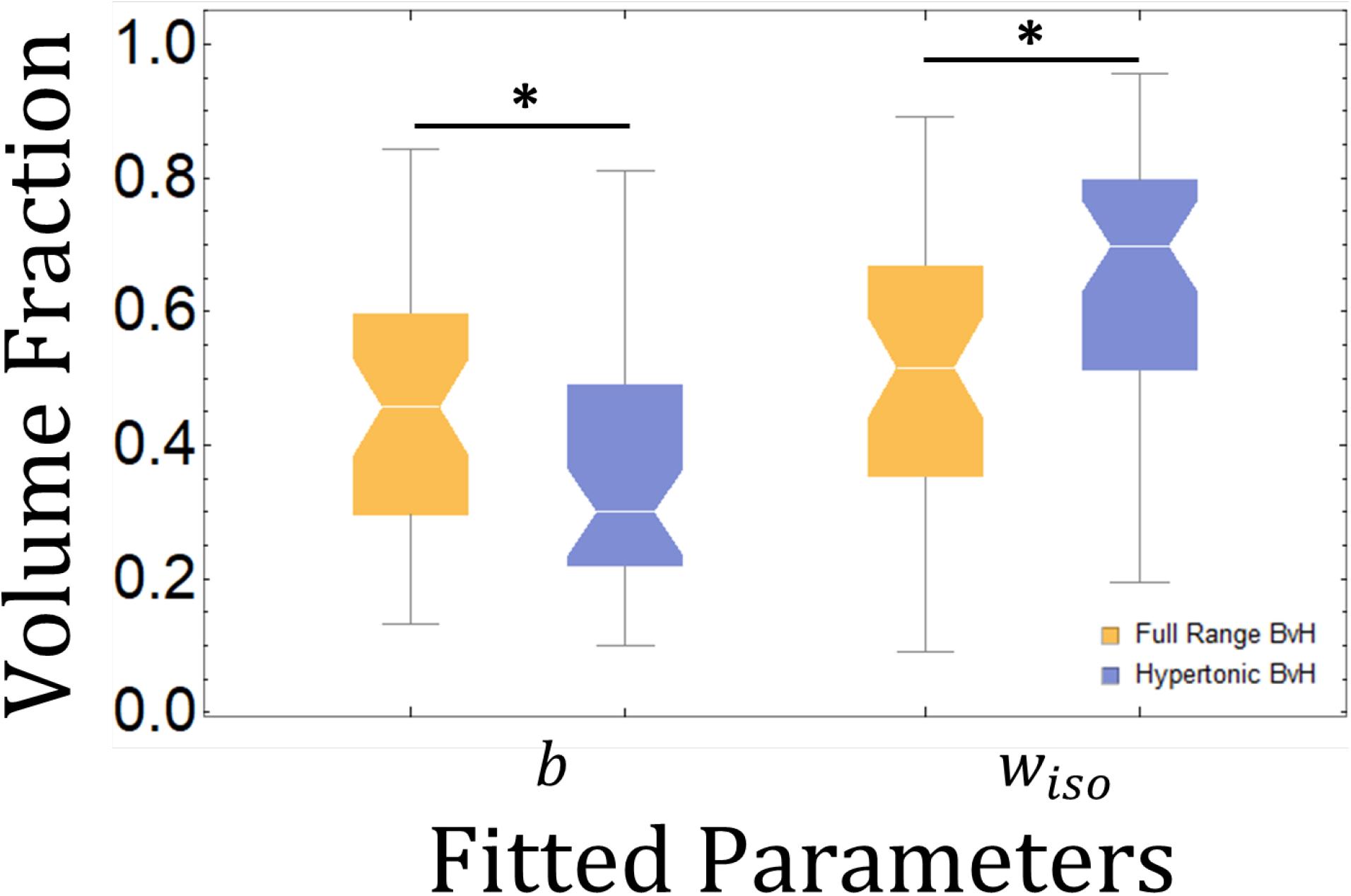
Group 3 notched plots for BvH relation fit (full range BvH versus hypertonic BvH) for 44 datasets. Notches indicated median. Stars indicate a significant difference of medians (*p*<0.05).

**Figure 5.**
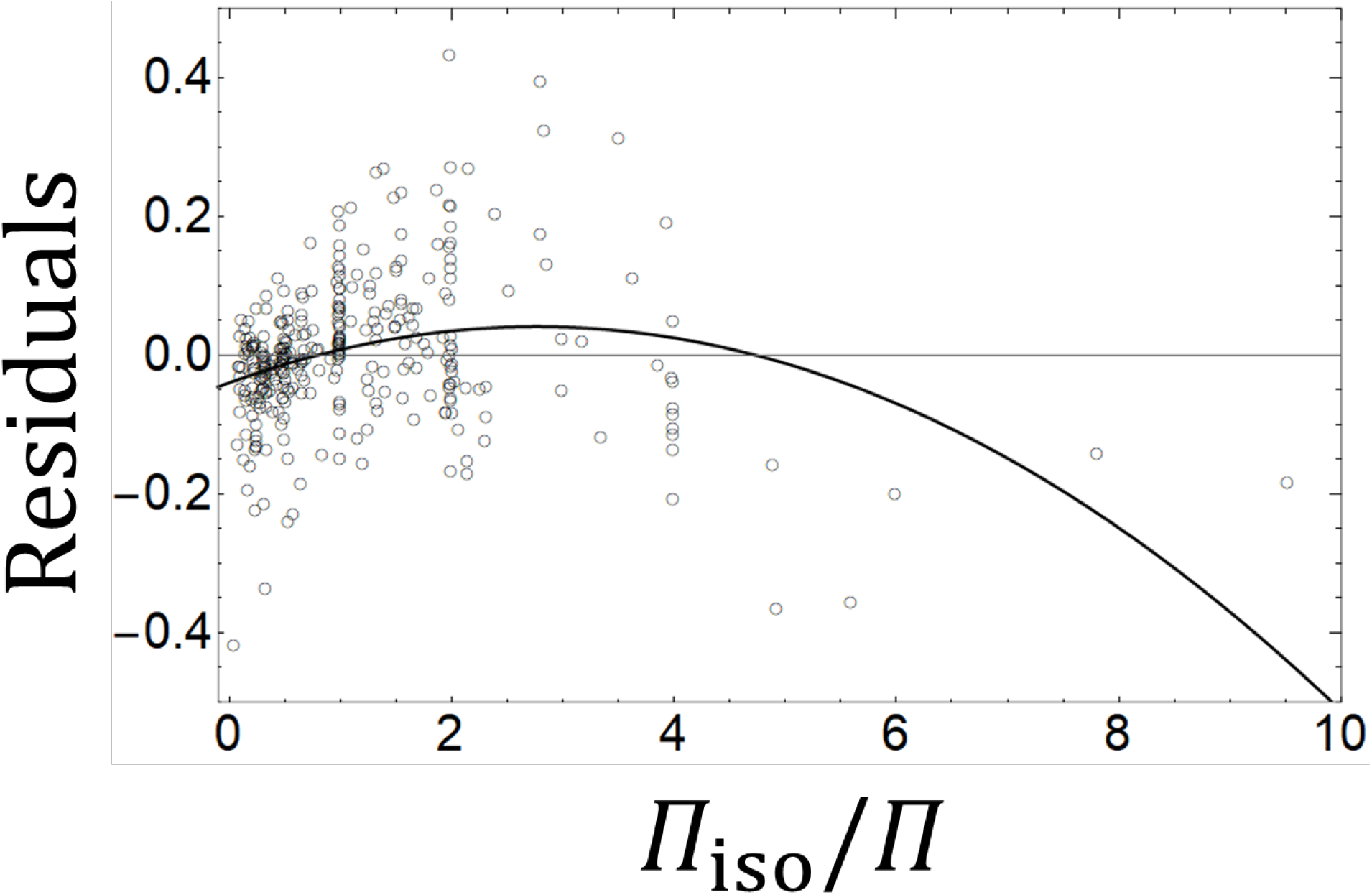
Fitted residuals of the BvH relation for Group 3 containing 44 datasets and 326 data points spanning the hyper- and hypotonic regions. The thick black line is a fitted second order polynomial providing a visual aid. For more information, see SM3.

**Figure 6.**
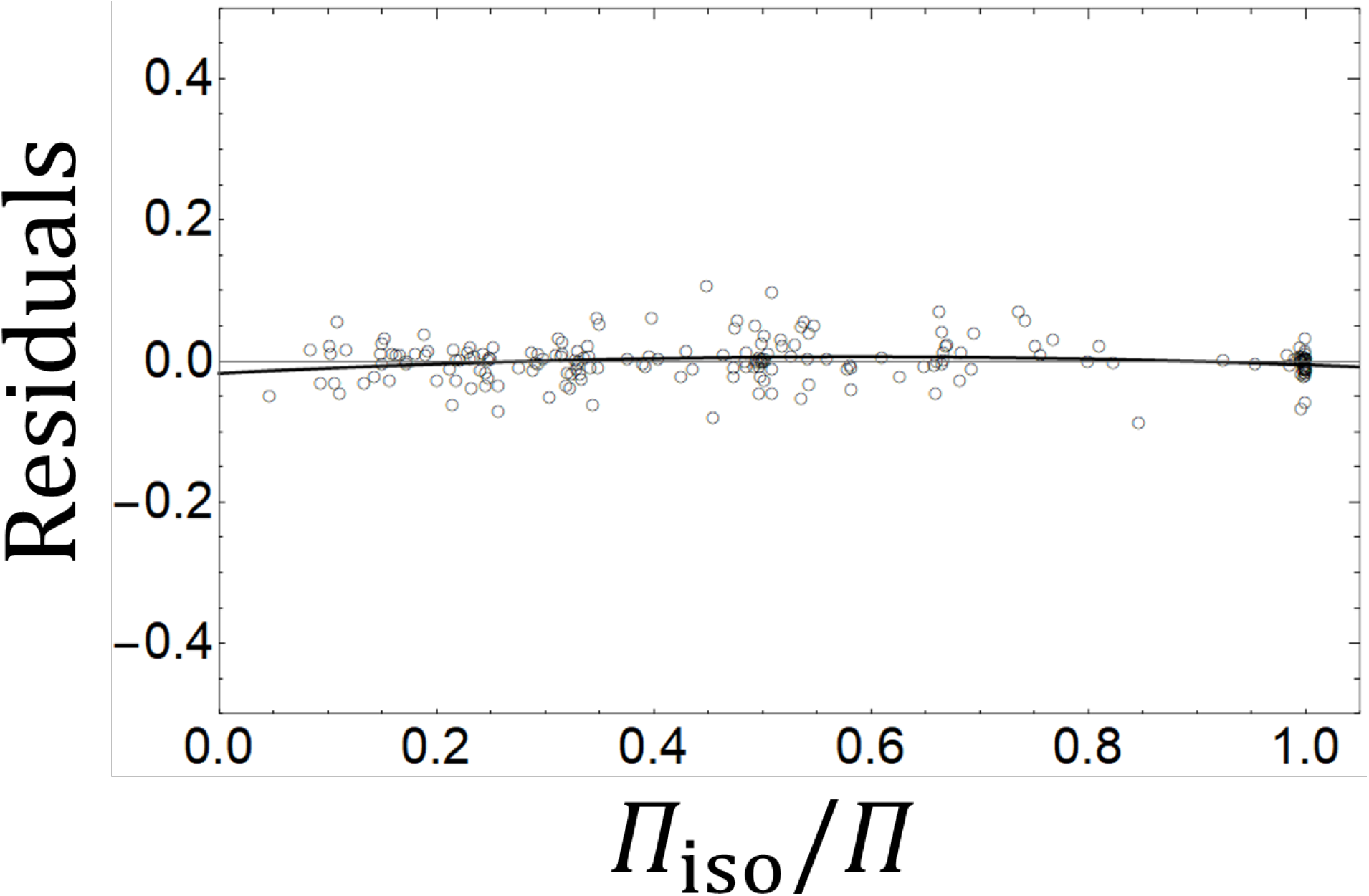
Fitted residuals of the hypertonic BvH relation for Group 3 containing 44 datasets and 202 data points in the hypertonic range. The thick black line is a fitted second order polynomial providing a visual aid. For more information, see SM3.

Along with the BvH relation, the leak model, and turgor model were fit to each of the 44 datasets of Group 3. The residuals of the leak model (Figure 7) and the turgor model (Figure 8) were plotted and neither had a noticeable trend in the residuals. The models are shown together fitted to each of the datasets in Group 3 (Figure 9). Comparisons between models are summarized in Table 1. To compare between nonlinear models, we found the total mean squared error (TMSE) for each dataset with the respective model. The turgor model had the lowest TMSE (0.099) closely followed by the leak model (0.105), and for reference, the linear BvH relation had the highest TMSE (0.478). To account for the number of parameters used, we computed the mean of the adjusted R^2^ value for each dataset with respect to each model. Both nonlinear models had very close mean adjusted R^2^ values with 0.9975 for the leak model and 0.9974 for the turgor model while the adjusted R^2^ medians (0.9980, and 0.9981 respectively) are not significantly different (Mann-Whitney U=908, *p*=0.6136). The nonlinear adjusted R^2^ cannot be directly compared to the linear BvH relation, which, for completeness, has a mean adjusted R^2^ of 0.9341, and median of 0.9647. Finally, to account for possible overfitting and small sample sizes, we ranked each model with respect to the computed AICc for each dataset, such that the model with the lowest AICc value was given one point (indicating the best performing model for the respective dataset). The total AICc score is then the number of times the respective model had the lowest AICc score for Group 3. The BvH relation had the highest ranked score of 33 while the turgor model had a score of 7 followed by the leak model with a score of 4. Notably, 16 of the BvH relation points were due to the nonlinear models being over fitted (given AICc of positive infinity) with respect to the number of data points. More information of each particular dataset is provided in the Supplementary Material (SM3).

**Figure 7.**
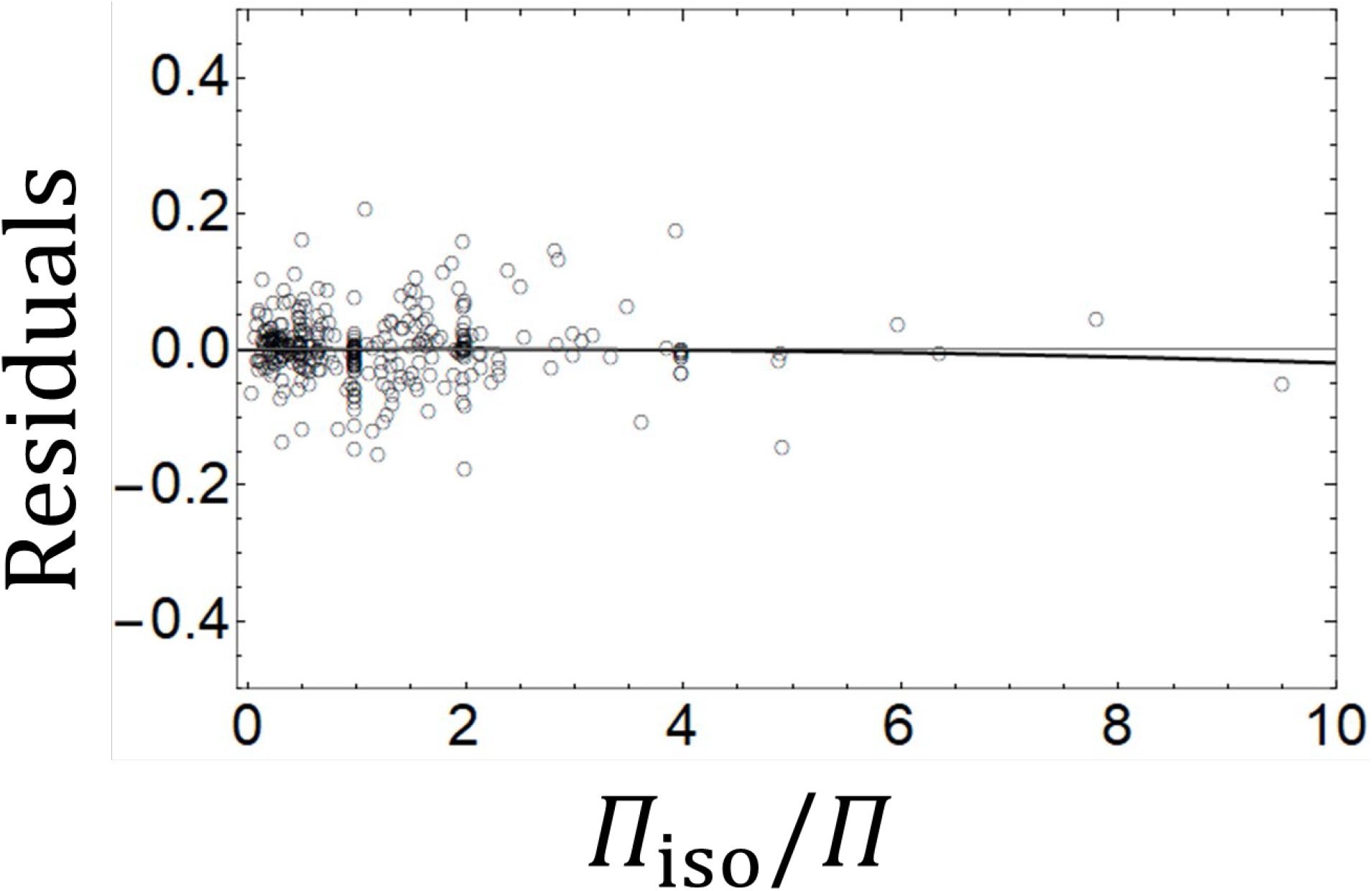
Fitted residuals of the leak model for Group 3 containing 44 datasets and 326 data points spanning the hyper- and hypotonic regions. The thick black line is a fitted second order polynomial providing a visual aid. For more information, see SM3.

**Figure 8.**
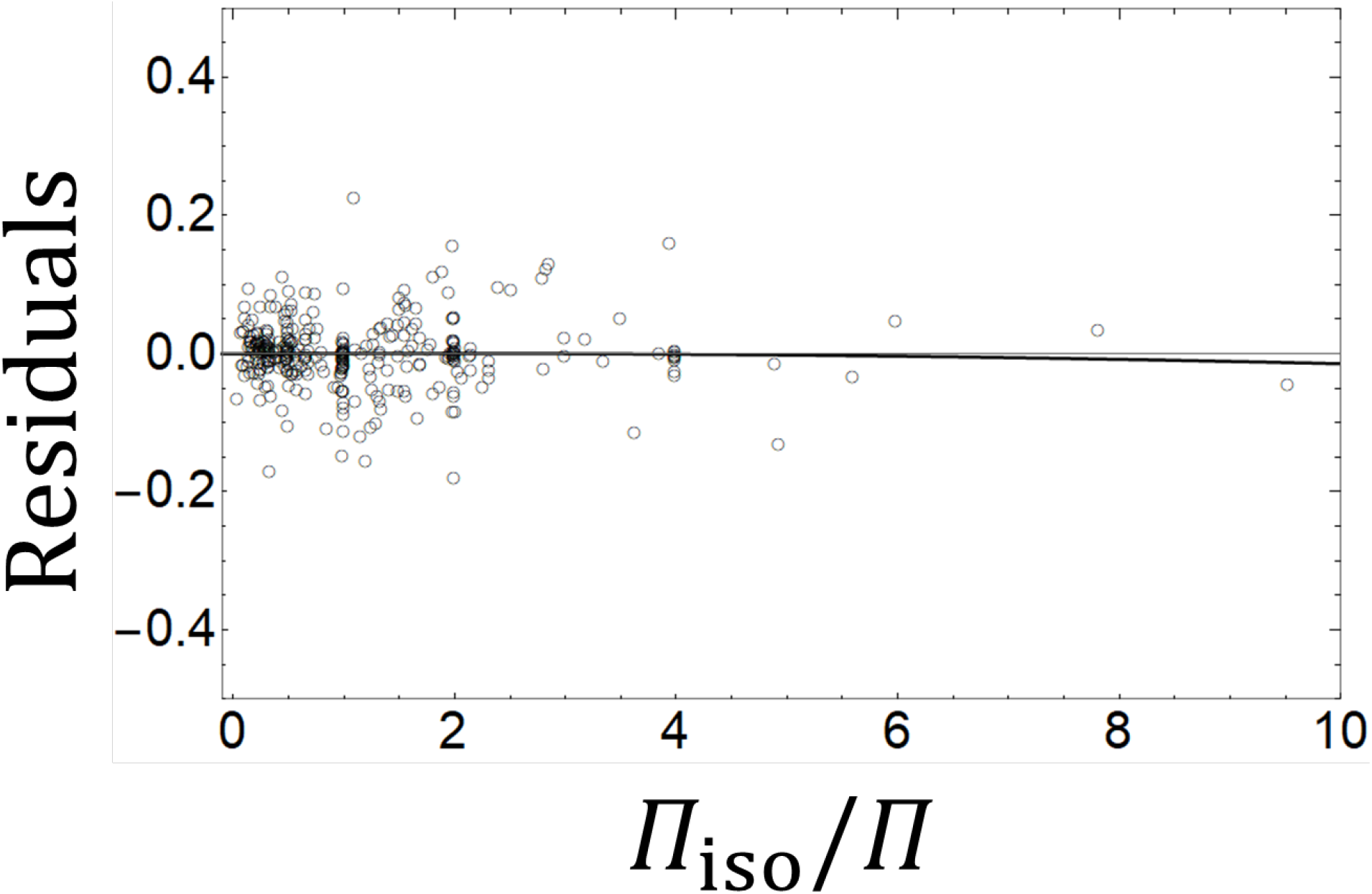
Fitted residuals of the turgor model for Group 3 containing 44 datasets and 326 data points spanning the hyper- and hypotonic regions. The thick black line is a fitted second order polynomial providing a visual aid. For more information, see SM3.

**Figure 9.**
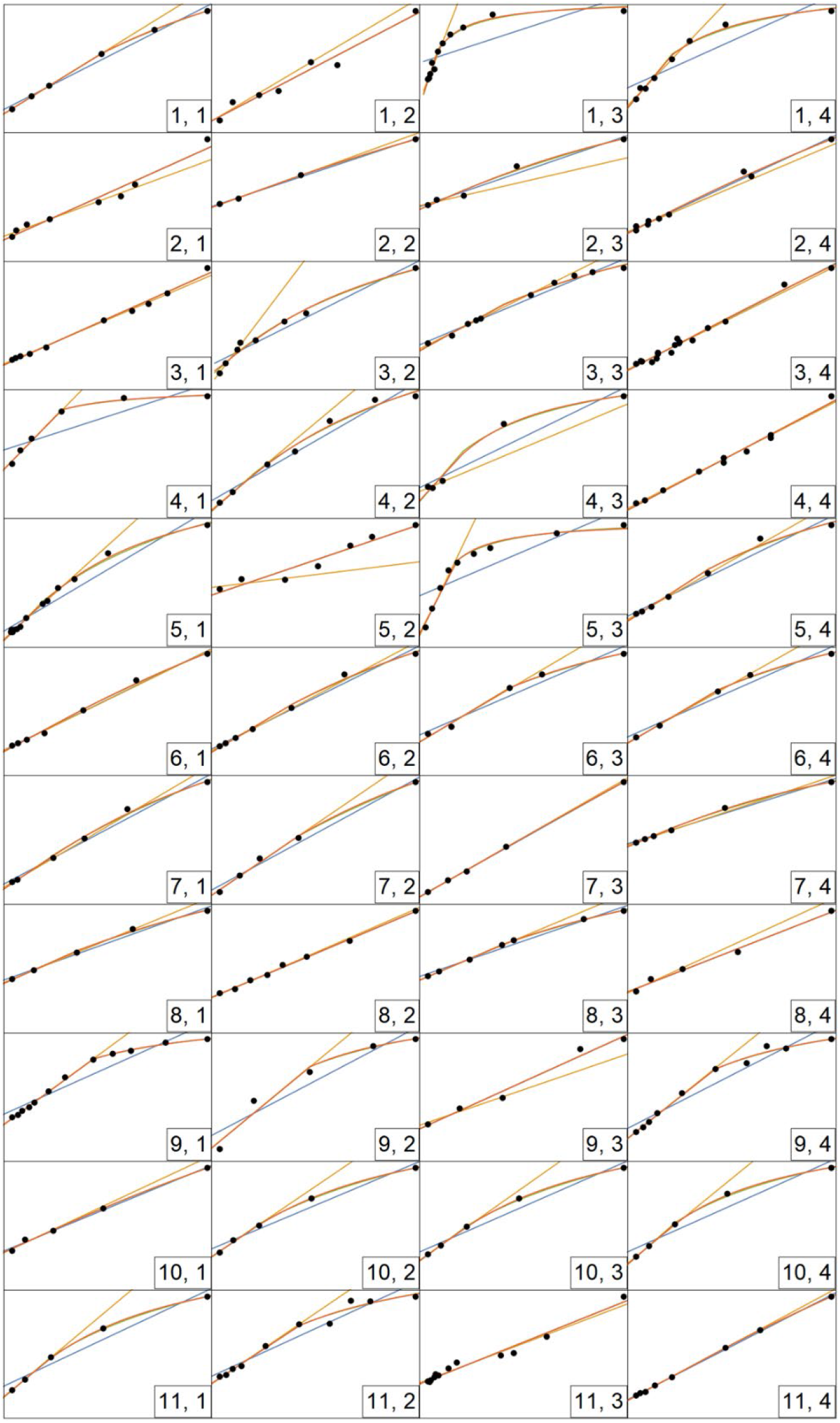
BvH plots of Group 3 with BvH relation (blue line), hypertonic BvH relation (orange line), leak model (red line), and turgor model (green line). Note the turgor model is almost never apparent due to being covered by the leak model. The vertical axes are relative cell volume (nondimensionalized) and the horizontal axes are in terms of inverse osmotic pressure (ranges vary among subplots). See SM3 for more information on each dataset.

**Table 1.** Summary of model comparisons for Group 3 containing 44 datasets with a total of 326 data points. Each model was fit independently to each dataset. The score for Ranked AICc is the total number of times the respective model has the lowest AICc out of all the tested models. Note the BvH relation had the lowest AICc score for 16 datasets due to low number of datapoints since the nonlinear models were considered overfitting and given the AICc score of positive infinity. See SM3 for more information.

| | # fitted<br>Parameters | Total Mean Squared<br>Error | (Mean, Median)<br>adjusted $R^2$ | Ranked AICc<br>Score |
| --- | --- | --- | --- | --- |
| BvH relation | 2 | 0.4782* | 0.9341*, 0.9647* | 33 |
| Leak Model | 3 | 0.1048 | 0.9974, 0.9980 | 4 |
| Turgor Model | 3 | 0.0989 | 0.9975, 0.9981 | 7 |

\* The Total Mean Squared Error and Mean, Median adjusted $R^2$ for BvH relation should not be directly compared to the nonlinear models as the BvH relation is a linear model and has one less parameter. The values are included here for completeness.

### 2.2 HepG2 Experiments

Using optical imaging, we obtained the BvH plot of HepG2 cells at room temperature (21 °C) and fit these data with the BvH relation and BvH relation restricted to the hypertonic region (Figure 10). The leak model and turgor model are fit together to the BvH plot and are nearly indistinguishable when plotted (Figure 12 A). The fitted parameters of each model are summarized in Table 2. The BvH relation is positively autocorrelated with a DW score of 1.466 which is lower than the critical bound (*n*=600, α=0.05) of 1.863. The residuals of the BvH relation have a distinctive downward bend (Figure 11 A), while this trend is absent for the residual plots of leak model (Figure 11 B) and the turgor model (Figure 11 C). The nondimensional BvH relation fitted *w*_iso_ (0.309±0.012 [SE]) is the lowest when compared to the leak model (0.845±0.053) and the turgor model (0.826±0.053). Reversely, the BvH relation fitted *b* (0.520±0.025) is larger when compared to the leak model (0.172±0.039) and the turgor model (0.182±0.039). The hypertonic BvH relation has very similar fitted values to the nonlinear models with a *w*_iso_ of 0.839±0.045 and *b* of 0.177±0.029. The leak model’s nonlinear term *λ* was fitted as 3.684÷0.600×10^−3^ while the turgor model’s nonlinear term *κ* is 2.382±0.411. Given the average cell radius is 7.42 μm and assuming the combined membrane and cortical thickness is 0.5 μm [4], then the fitted young’s modulus is approximately 6.27 MPa.

**Figure 10.**
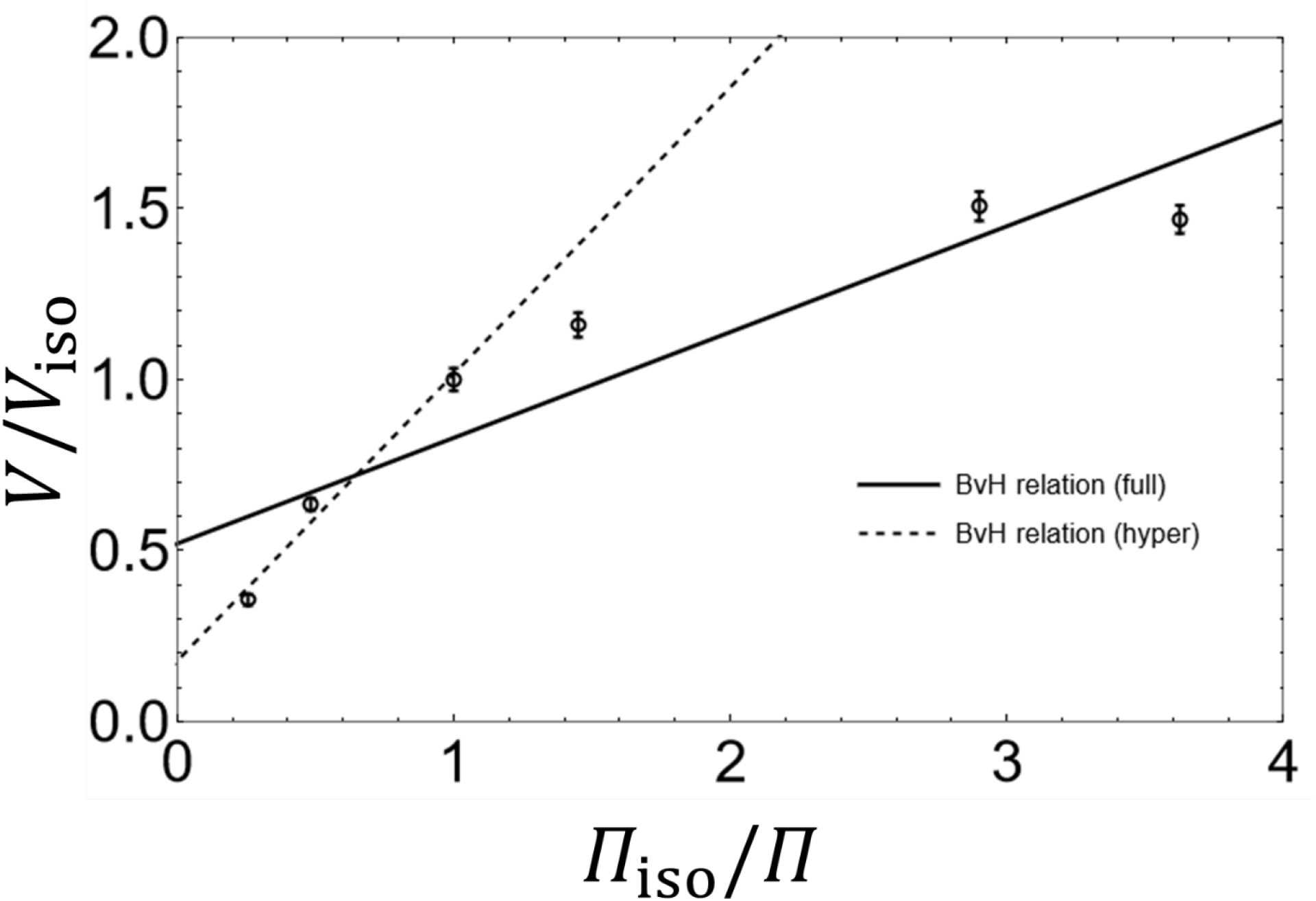
HepG2 BvH plot at room temperature (*n*=600). Error bars are standard error of the mean.

**Figure 11.**
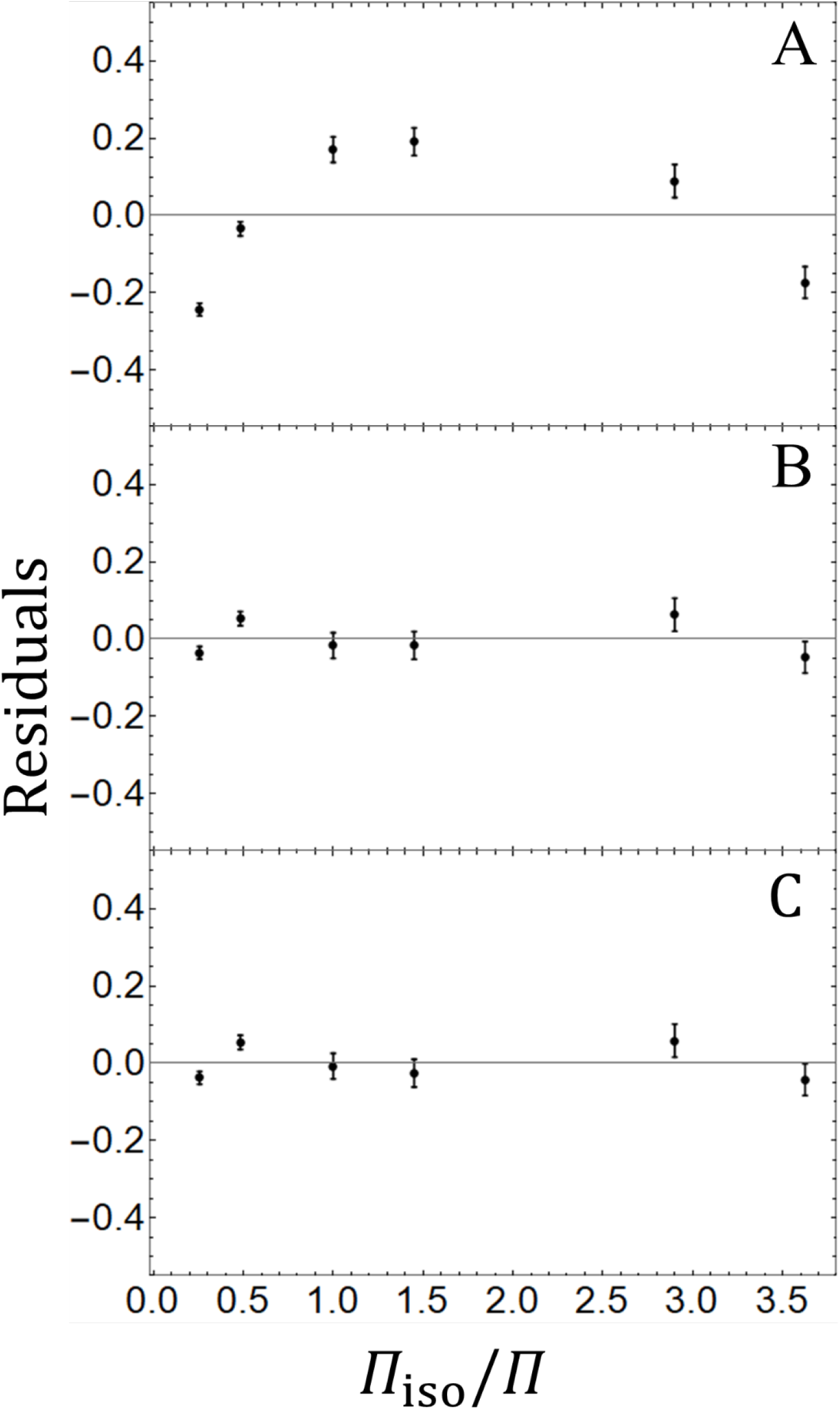
Residual plot for (**A**) BvH relation, (**B**) leak model, and (**C**) turgor model of HepG2 cells (*n*=600). Error bars are standard error.

**Figure 12.**
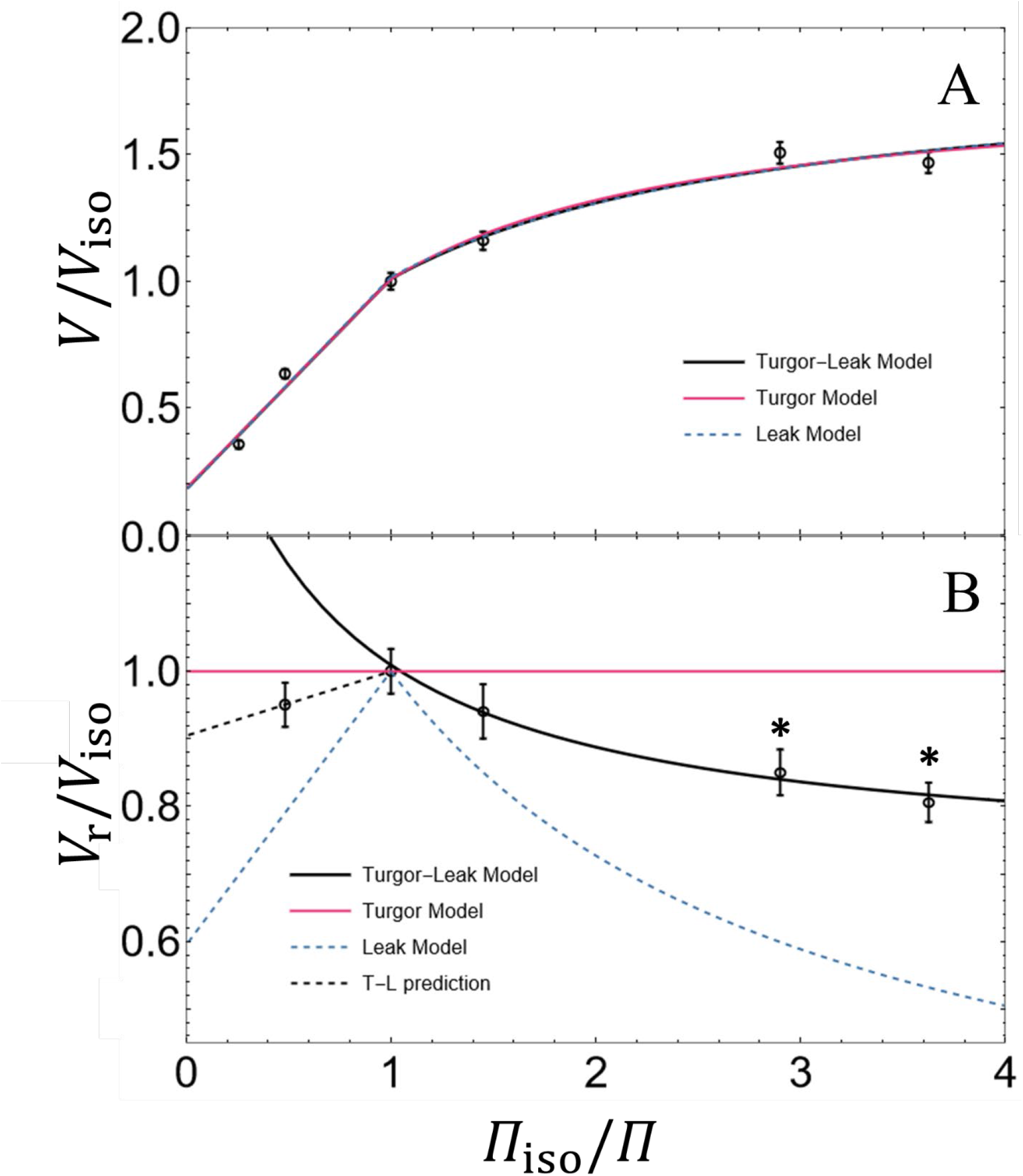
Turgor-leak model fit (*Eq*. 7: solid black line) to the HepG2 BvH plot (**A**: *n*=600) and return to isotonic conditions plot (**B:***n*=400), with the hypertonic point not being included in fitting (*n*=100). T-L prediction (dashed black line) is the predicted return to isotonic volume using *Eq*. 5 with the estimated *λ* value from *Eq*. 7. Error bars represent standard error. Stars indicate significant difference (*p*<0.05) between the respective point and the isotonic point.

**Table 2.** Fitted parameters for HepG2 modelling. Fitted parameters are followed by ± SE.

| | $w_{\text{iso}}$ | $b$ | Nonlinear constant(s) | $n$ |
| --- | --- | --- | --- | --- |
| BvH relation | 0.309±0.012 | 0.520±0.025 | | 600 |
| BvH relation (hyper) | 0.839±0.045 | 0.177±0.029 | | 300 |
| Leak Model | 0.845±0.053 | 0.172±0.039 | $\lambda = 3.68 \pm 0.060 \times 10^{-3}$ (unitless) | 600 |
| Turgor Model | 0.826±0.053 | 0.182±0.039 | $\kappa = 2.38 \pm 0.392$ (unitless) | 600 |
| Turgor-Leak Model | 0.830±0.047 | 0.179±0.038 | $P_n = 5.54 \pm 1.095 \times 10^{-8}$ dm s<sup>-1</sup><br>$K = 3.93 \pm 0.605$ MPa | 1000 |

To help differentiate between the two nonlinear models, we plotted volume after return to isotonic conditions as a function of anisotonic challenge along with the predicted return volumes of the leak model (*Eq*. 5) and the turgor model (no change from initial isotonic volume) (Figure 12 B). An ANOVA indicated there was a significant difference between return volumes (F_[4,495]_=5.419, *p*=2.82×10^−4^) while a post-hoc Bonferroni test verified cells equilibrated to 100 mOsm/kg and 80 mOsm/kg have a significantly lower return volume than the isotonic volume (*p*<0.05). A T-Test indicated both of these points were significantly different (*p*<0.05) from the leak model prediction (*Eq*. 6).

We simultaneously fit the turgor-leak model (*Eq*. 7) to both the BvH plot (Figure 12 A) and the return to isotonic conditions plot (Figure 12 B). Using a hydraulic conductivity parameter *L*_p_ of 1.094 dm s^−1^ kPa^−1^ [49], our fitted *P_n_* was 47.6±9.11×10^−9^ dm s^−1^, and the Young’s modulus was fit to 3.85±0.44 MPa. The turgor-leak model is nearly indistinguishable from the other nonlinear models in the BvH plot but differs significantly in the return to isotonic conditions plot. Fitting the turgor model and leak model to the BvH plot and return to isotonic plot simultaneously, provides a total fitted AIC and BIC for each model, enabling a direct comparison to the turgor-leak model. The turgor-leak model has the lowest AIC and BIC values out of all the nonlinear models (Table 3). The analogues *λ* value for the turgor-leak model was 4.82×10^−4^, when *λ* = *ρ*_s_*P_n_*/*RTL*_p_ and the ion molar volume, *ρ*_s_, is 0.027 L mol^−1^ [18]. Using the turgor-leak derived *λ* value, *Eq*. 5 better predicts the hypertonic (600 mOsm/kg) return to isotonic volume than *Eq*. 7 (Figure 12 B).

**Table 3.** Comparison between nonlinear models fitted to BvH plot and return to isotonic conditions plot (*n*=1000).

|  | Num. Parameters | AIC | BIC |
| --- | --- | --- | --- |
| Leak Model | 3 | 704.85 | 724.48 |
| Turgor Model | 3 | 712.15 | 731.78 |
| Turgor-Leak Model | 4 | 659.47 | 684.01 |

## 3 Discussion

### 3.1 Meta-analysis

First, the results of the meta-analysis support the hypothesis that the BvH relation cannot be an *a priori* assumption about cell osmotic response. A significantly larger number of datasets (6 out of 28; 21.4%) were autocorrelated using a linear fit than expected (5%). These statistical findings are supported by the downwards bend in the residuals plot for the BvH relation when regressed with respect to both hyper- and hypotonic regions (Figure 2). A downward bend in the residuals was again found for the larger dataset of Group 3 (Figure 5). This indicates that a linear model is not appropriate in general and by extension not all cells are governed by the BvH relation. Indeed, some cells appear to have a linear relation while others are highly nonlinear (Figure 9). Therefore, models predicting cellular volumetric response to osmotic conditions must be determined at the cell specific level.

Next, the results of the meta-analysis do not reject the hypothesis that the BvH relation can be assumed for cells *a priori* in hypertonic conditions. There was not a significantly larger number of hypertonic datasets (2 out of 22; 9.1%) autocorrelated when linearly regressed than expected (5%). There is no apparent general trend in the residuals for the BvH relation fitted to only the hypertonic region of Group 2 (Figure 3). There was no residual pattern found for the larger dataset of Group 3 (Figure 6). However, the fitted BvH relation to the hypertonic region may not necessarily be applicable to the respective cell when crossing to the hypotonic region. Fitting a piecewise linear model may be a more appropriate first order approximation of equilibrium volumes, but does not necessitate that the fitted parameters have physiological importance [30,41].

Failing to reject the BvH relation for the hypertonic region as a “good model” does not dispute the existence of regulatory volume increase (RVI) but instead indicates the utility of the simplistic BvH relation for cells in general in the hypertonic region. Furthermore, RVI is known to be much slower than regulatory volume decrease (RVD), and therefore changes in volume due to RVI may simply be insignificant during short time periods associated with the BvH plot [1,54,55]. A more complex RVI model should only be used for a cell which is known to undergo RVI. In such a case, a more complex model may be warranted that may require more experimentation than required for a BvH plot.

Prickett and colleagues [32] identified a non-ideal replacement equation for the BvH relation for non-ideal non-dilute solutions and apply this model for erythrocytes. For our collected data the effect of non-ideal solutions was not significant enough to rebuke the BvH relation in the hypertonic region for cells in general as there were not a significant number of autocorrelated datasets and no obvious trend in the fitted residuals (Figure 3, Figure 6). The majority of the mean data points collected in the hypertonic region were in the range 0.1 to 1.0 (e.g. 3000 mOsm to 300 mOsm) and therefore are not necessarily as “non-ideal” as more concentrated solutions (depending on the solution). In this 0.1 to 1.0 hypertonic range, the assumption that osmolality is proportional to molality may hold for many solutions [26,56]. It is possible that an autocorrelated trend would appear for higher osmolalities but no such evidence is found in the present data. If the non-ideal non-dilute BvH relation is generally applicable for cells, then further research must be undertaken for multiple cell types across the so-called “non-ideal” region, that is, the region wherein intracellular osmolality is not proportional to molality.

The change in accuracy of the BvH relation over the hyper- and hyptonic regions versus just the hypertonic region is directly reflected in the fitted *w*_iso_, and *b* parameters (Figure 4). The fitted non-osmotic volume is lower when fitting to just hypertonic region as opposed to fitting to both the hyper- and hypotonic region, and the reverse is true for osmotically active water volume (Figure 4). Both nonlinear models tend to agree with the hypertonic fit of the BvH relation, in part because of the nonlinear models’ piecewise structure, but also because they account for this downwards bend which generally appears in the transition from hyper- to hypotonic region in the BvH plots (Figure 9). The claim that a cell behaves linearly as an ideal osmometer at the expense of omitting data or ignoring signs of autocorrelation may lead to inability to predict volumetric response and poorly informed physiological models [57,58]. Not accounting for the osmotic regulation of the cell may have direct consequences in the fitted osmotically active volume and non-osmotic volume. Fitting to just the hypertonic region may improve these fits but does not inform on the presence or absence of osmoregulation. Indeed, incorrectly assuming a cell is an ideal osmometer can result in poorly predicting volume during and after anisosmotic challenge as well as volumetric response at different temperatures [29,55,59,60].

The data points collected are mean data points (no variance collected). As such the fitted models are only with respect to the means of data points in the respective articles. The analysis is done in terms of checking the general behaviour of models and comparison between models, not in an effort to reobtain literature values. Our presented fitted parameters, adjusted R^2^, MSE, and AICc values for each dataset then are distinct from the values presented in their respective papers (if presented). Our fitted parameters then should not be considered a library of cell osmotically active volume and non-osmotic volume, but instead an estimate given the mean data points. Comparing the nonlinear models in terms of specific numbers (Table 1) must be taken in context of the fitted data (Figure 9) especially when comparing adjusted R^2^ values [61]. In terms of mean adjusted R^2^ values, the nonlinear models (leak: 0.9974, turgor: 0.9975) perform about equally well, with similar values in total mean squared error (leak: 0.1048, turgor: 0.0989). These similarities are reflected in how the models are almost always indistinguishable when plot (Figure 9). The ranked AICc score comparison indicates that while a majority of cell types are robustly predicted with a linear model, 11 out of 44 (25%) cell types are robustly predicted by nonlinear models (Table 1). Note that due to low number of data points for some datasets, the BvH relation had the lowest AICc score for 16 datasets since the nonlinear models were considered overfitting (e.g. fitting a 3 parameter model to 4 or 5 data points). Both the leak model and the turgor model perform near identically when plotted (Figure 9) despite being based on distinct mechanisms while neither have trends in their residual plots (Figure 7, Figure 8). Our results support the hypothesis that there is no one replacement general model for the BvH relation, instead there are cell specific models that may be identified based upon the underlying mechanism of osmoregulation (or lack thereof).

The underlying mechanism of osmoregulation is not necessarily known for each respective cell (if present). However, it is interesting to note how observed mechanisms for specific cell types match up with how well a specific model (that assumes those mechanisms) fits the respective data. The leak model notably excels at predicting the osmotic behavior of HeLa cells (dataset {9,1}), Chondrocytes (dataset {1,3}), and kidney fibroblasts (dataset {9,1}), which are thought to use mechanosensitive gates to release osmolytes [54,59,62]. Interestingly, some chondrocytes datasets appear linear (datasets {2,4},{8,2}) while others were very nonlinear (dataset {1,3}) potentially due to species differences but also the range of osmotic challenge. For cells with nonlinear behaving BvH plots, our fitted *λ* ranged from approximately 1.0×10^−3^ to 1.1×10^−2^. To our knowledge, Casula and colleagues [18] were the first to present the BvH nonlinear constant *λ* (based on their fitted *L*_p_ and *P*_n_ values at room temperature, *λ* is calculated to be 2.53×10^−6^). However, for each dataset the leak model accurately fits, the turgor model has an almost identical fit.

The membrane-cortex elastic modulus varies widely dependent on cell type. The modulus (typically in terms of Young’s Modulus) varies from 1 to 50 kPa of human platelets [63], 13 to 150 kPa for lung carcinoma cells [64], 670 kPa for rat atrial myocytes [65], 4.3 MPa for rat mast cells [66], and 10 MPa for human red blood cells [67]. Liu and colleagues [68] suggest the nonlinear behaviour of pancreatic islet cells are due to possible ionic leakage or mechanical resistance due to high abundance of filaments (see also [29,69,70]). Using the membrane-cortex thickness of 0.5 μm for animal cells suggested by Jiang and Sun [4], these pancreatic islet cells had a fitted Young’s modulus ranging from approximately 12.4 to 266 MPa (datasets {1,1}, {4,1}, {10,2}, {10,3}, {10,4}, {11,1}). These values are generally higher than previously recorded for other cells. On the other hand, our fittedλ values for pancreatic islet cells range from 9.62×10^−4^ to 1.29×10^−2^, approximately 2 to 4 magnitudes larger than presented by Casula and Colleagues [18]. A realistic range of *λ* is currently unclear, however, using both the mechanical resistance and ion leakage may reduce both fitted nonlinear values to a lower (and perhaps more reasonable) range.

As for the zona pellucida of oocytes, our fitted Young’s modulus for bovine MII zona pellucida (ZP) was 2.64 MPa (Figure 9: dataset {9,4}) using a ZP thickness of 10 μm [71]. This value is approximately an order of magnitude larger than estimates using atomic force microscopy (~100 to 600 kPa) [71]. Furthermore, our fitted Young’s modulus for mouse fertilized ova was approximately 197 kPa (ZP thickness of 4.5 μm reported by Sun and colleagues), which is approximately 5 times larger than recorded by point depression (Youngs modulus of 42.2 kPa) [72]. Both previous estimates of Young’s modulus are derived from local compression/manipulation methods and includes sheer stresses, while our estimates are derived from uniform swelling of a thin shell. The difference then is our estimates are due to in-plane deformation with little to no sheer stress, while local compression results in an out-of-plane deformation. Indeed, wire mesh materials have an effective in-plane elastic modulus of about an order of magnitude higher than out-of-plane modulus [73]. This difference in structural loading may be the key reason for differences in observed Young’s modulus.

Simply fitting the turgor model or leak model to a BvH plot has been shown in this paper to have effectively the same shape and overlap for all nonlinear models (Figure 9), without a method of differentiation, one cannot exclude the possibility of either mechanism. Supplementation of the BvH plot with a return to isotonic conditions plot tests the hypothesis of purely mechanical resistance (predicted return to isotonic volume is the same as initial isotonic volume), or purely leak driven (predicted return to isotonic volume is lower than initial isotonic volume and follows *Eq*. 5). However, one can also obtain a combination of both mechanical resistance and leakage from the combination of both plots using *Eq*. 7. We note that this methodology is improved by taking time dependent measurements of cells throughout time to reduce overfitting when fitting for *w*_iso_, *b*, *L*_p_, *P*_n_, and *K*. Fitting at both physiological (e.g. 37 °C), room temperature, and cold temperatures (e.g. 4 °C) [1,41] may further improve this model by enabling an estimate of active pumping importance for the particular cell given *Eq*. 7 is extended across temperatures via the Arrhenius equation [11].

### 3.2 HepG2 Experiments

Our results show that HepG2 cells are significantly autocorrelated with respect to the BvH relation and are therefore non-ideal osmometers (Figure 10). Both nonlinear models perform better than the BvH relation but appear to overlap each other for the majority of the BvH plot (Figure 12 A). The turgor model’s fitted Young’s modulus of 6.27 MPa is roughly comparable to the Young’s Modulus of 4.3 MPa for rat mast cells [66]. Using the leak model, the fitted *λ* (3.68×10^−3^) value is about 3 orders of magnitude larger than that fitted by Casula and colleagues (2.53×10^−6^) [18]. Since the turgor model assumes no ionic leakage while the leak model does, the return to isotonic volumes plot is used to help differentiate the two models (Figure 12 B). The turgor model is ruled out as being the sole mechanism for osmoregulation as return to isotonic volumes are significantly different from initial volumes. The leak model prediction is significantly (*p*<0.05) smaller than observed return isotonic volumes indicating ion leakage is not the sole mechanism of osmoregulation.

By using both the BvH plot and the return to isotonic conditions plot (Figure 12), we fit the turgor-leak model (*Eq*. 7) and found lower fitted values for both the Young’s Modulus (3.85 MPa) and *λ* (4.82×10^−4^) in comparison to the turgor model and leak model respectively. When fit to both BvH plot and return to isotonic conditions plot, the turgor-leak model has a lower AIC and BIC when compared to either the turgor model or leak model (Table 3). This indicates the turgor-leak model is the most accurate model presented despite having one more fitted parameter. We do note that if literature value of L_p_ were not accessible then fitting *Eq*. 7 would likely require inclusion of time dependent data to avoid overfitting.

Interestingly, using the turgor-leak *λ* value with *Eq*. 5 (the leak model return to isotonic conditions), the predicted return to isotonic volumes after hypertonic challenge is greatly improved (Figure 12). The difference is that the turgor-leak model assumes the membrane-cortex does not change structure over time but instead passively forces water into the cell along with ions, meaning the cell will have more ions inside and thus a higher return to isotonic volume. On the other hand, the leak model assumes the membrane-cortex self regulates, changing its structure to accommodate its smaller size, such that when returning to isotonic conditions the cell swells, activates mechanosensitive ion channels, releases ions, resulting in a lower return to isotonic volume. Future work may investigate the effects of cytochalasin D, temperature, and ion channel blockers on mechanobiological dynamics especially in consideration across hyper- and hypotonic regions. Such efforts will improve our understanding of osmoregulation and by extension dynamic volumetric responses of cells.

## 4 Materials & Methods

### 4.1 Models

#### 4.1.1 Non-Ideal Non-Dilute Model

Prickett and colleagues provide a derivation of a model similar to the BvH relation in terms of conservation of mass and correct the BvH relation for non-ideal non-dilute (NIND) solutions [32,74,75]. The moles of non-permeating solute *N*_s_ inside the cell remains constant such that,

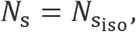

or,

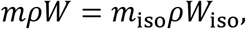

where *m* is the molality of intracellular solutes, *ρ* is the density of water, *W* is the solvent water volume of the cell, and the subscript iso is the respective quantity at the isotonic point for the cell. For a given osmotic condition, relative cell volume can be written as a function of molality in the form,

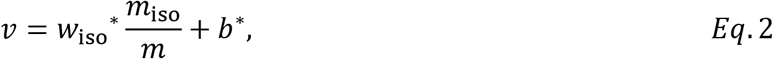

where *w*_iso_^*^, and *b*^*^, are the osmotically active fraction and non-osmotic fraction, respectively, without the ideal dilute assumption. Intracellular and extracellular osmolalities are equal at equilibrium, this requires the intracellular molality to be written in terms of osmolalilty which can be done by using the osmotic virial equation *π*(*m*), and its inverse *m*(*π*):= *π*^−1^(*m*) [32,74,75]. For many solutes such as NaCl the van ’t Hoff factor *i* provides a good estimate for a range of concentrations that are physiologically relevant (e.g. < 1 mol/kg) [75]. Some proteins and solutes such as hemoglobin and dimethyl sulfoxide behave non-ideally and as such the respective virial equation takes the form of a polynomial such that *π*(*m*) = *A m* + *B m*^2^ + *Cm*^3^, where *A*, *B*, and *C* are fitted constants [32,75]. It is this nonlinear relation between osmolality and molality that results in a nonlinear downwards bend in the extreme hypertonic regions of a BvH response [75], reducing predicted non-osmotic fraction of a cell when compared to the BvH relation [32]. This can have serious implications for mass transfer modeling and equilibration volumes. However, Elmoazzen and colleagues [56] argue that without an abundance of large intracellular proteins, a cell containing solutes mostly composed of salts will have osmolality proportional to molality for a wide range of osmotic conditions, which is echoed by Benson [26]. In these cases, the predicted non-osmotic fraction of cells using the BvH relation (*Eq*. 1) and the NIND model (*Eq*. 2) will be very close. To date, there have been few efforts to determine if the majority of cells in general are linear in the sufficiently non-dilute hypertonic region, thus it remains unclear if a linear approximation for *π*(*m*) is appropriate for most cells, or that a linear model is insufficient for cells in the hypertonic region [26,56].

#### 4.1.2 Leak Model

One method for cells to reduce catastrophic damage under large osmotic gradients is to change their lipid membrane permeability to intracellular osmolytes [60]. Cells with anionic non-permeable molecules (e.g. proteins and organic phosphates) are threatened with continuous cell swelling due to diffusion of ions and concomitantly water into the cell. Since membranes are relatively impermeable to ions, this swelling can be mitigated via actively pumping ions across the membrane [13–15]. This pumping of ions is accompanied by a facilitated diffusion of Cl^−^ ions and is collectively referred to as a pump and leak concept [1,15]. As ionic diffusion across the membrane is several orders of magnitude lower than water, the amount of ions needed to be actively pumped to maintain dynamic equilibrium is low. The pump and leak concept captures the ability for cells to both actively and passively regulate intracellular ions to reach both a membrane potential equilibrium and an osmotic equilibrium [1,4,15]. However, on short time scales (minutes to tens of minutes) of high osmotic gradients, active pumps may not significantly influence the osmotic response and instead osmotic response is dependent on facilitated diffusion.

By selectively and possibly temporarily changing the cell membrane to become permeable to some osmolytes, the cell allows osmolytes to leave or possibly enter during anisosmotic challenges [54,59,76,77]. To do so, one method of changing membrane permeability is by opening or closing mechanosensitive channels embedded on the cell membrane [18,78]. Shrinking or swelling of the cell results in a change of membrane curvature and membrane tension, which may activate associated channels enabling the cell to respond to osmotic stresses [79–81]. A leak model takes into account the cell membrane as an active and dynamically structure, changing in area and molecular components to respond to environmental changes [60,80,82,83].

Both the BvH relation and the NIND model assume a constant amount of intracellular solute molecules. During hypotonic swelling, a loss of intracellular molecules across the membrane will lead to a smaller equilibration volume than predicted by these models. If the cell gains solutes while shrinking in hypertonic conditions, then the equilibration volume will be larger than predicted by the above models. We provide a simple predictive model as follows. According to the BvH relation, *inRT*/*ρ* is constant. If *Π*_iso_*W*_iso_ = *inRT*/*ρ*, but *Π*_hypo_*W*_hypo_ ≠ *inRT*/*ρ*, or *Π*_hyper_*W*_hyper_ ≠ *inRT*/*ρ*, then it is possible that intracellular solute molecules have left or entered the cell respectively. The new possible intracellular solute amounts results in new values, namely *in*_hypo_*RT*/*ρ*, and *in*_hyper_*RT*/*ρ*. These new values have predictive associated return to isotonic water volumes, *W*_r_, where in particular,

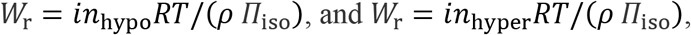

in hypotonic and hypertonic conditions, respectively.

These predictions assume *in*_hypo_*RT*/*ρ* and *in*_hyper_*RT*/*ρ* remain constant while returning to isotonic conditions. If cells appear nonlinear in the BvH plot, then the difference between the expected volume and the observed volume will provide the expected final return volume, *V*_r_ = *W*_r_ +*V*_b_, which then can be verified directly. Assuming a linear hypertonic region, this model can be written in terms of observed volume during anisotonic challenge, *V*_obs_, in the form,

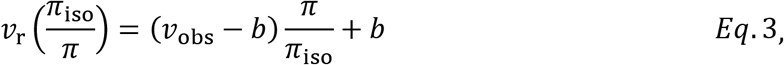

where *ν*_r_ = *V*_r_/*V*_iso_, and *ν*_obs_ = *V*_obs_/*V*_so_, and *b* is fitted via the BvH relation in the hypertonic region assuming a linear relation. If a linear hypertonic relation is not present, then *b* may be fit to data from the BvH plot the return to isotonic conditions plot simultaneously by letting *ν*_r_ be a function of both *ν*_obs_ and *π*_iso_/*π*.

A more complex leak model was developed by Casula and colleagues [18]. This model accounts for changes in cell membrane area as well as ionic leakage in determining volumetric equilibrium with respect to anisosmotic conditions. This “leak model” includes mechanosensitive channels that become permeable to internal ions during cell swelling. In particular, Casula and colleagues [18] argue that when the tension in the cell membrane is above a certain threshold, then mechanosensitive channels are open and ions may flow down their respective concentration gradients. Their model uses four parameters, and thus to minimize overfitting (e.g. fitting 4 parameters to 5 to 10 mean data points), in this manuscript we set the threshold tension to the isotonic tension such that any swelling past isotonic volume results in opening of these channels. This is equivalent to assuming that solute molecules are lost during swelling in only hypotonic regimes. The model results in linear behavior while shrinking and a nonlinear behavior while swelling. Assuming the cell starts from isotonic volume and then exposed to anisotonic conditions, the leak model takes the form,

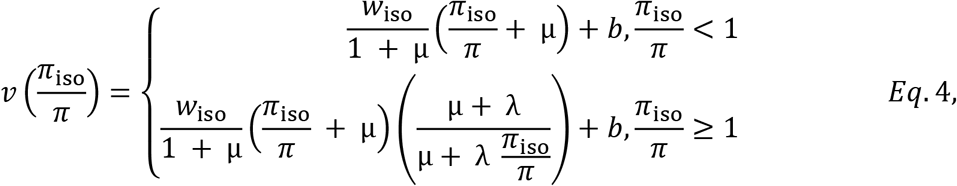

where *ν*, *w*_iso_, *b*, and *π*_iso_/*π*, are described as in *Eq*. 1, μ = *ρ*_s_ *π*_iso_/*s* where *p*_s_ is the molar density of salts, s is dissociation of salts, and *λ* = *P*_n_/(*R T L*_p_) where *P*_n_ is permeability of ions across the cell membrane, *L*_p_ is hydraulic conductivity of the plasma membrane. Our *Eq*. 4 differs from Casula and colleagues [18] by the explicit inclusion of *w*_iso_ which must be fit when isotonic volume is not known for each sample [26]. Note that if *λ* is zero then the model is equal to the BvH relation [18]. Furthermore, since *π*_iso_/*π* ≅ *c*_iso_/*c* in dilute solutions, this particular volume equilibrium model may be used with both osmolality and osmolarity in the hypotonic region, with the predictive volume error between the two being <1% for a common solution such as 1x PBS (Phosphate Buffered Saline at 300 mOsm/kg), though the models may have different fitted parameters.

The model presented by Casula and colleagues [18] may also be written to predict return to isotonic volume after osmotic challenge. The change in volume after return to isotonic conditions is a function of loss of intracellular ions, which in turn is a function of the duration the mechanosensitive channels are open. The return relative cell volume, *ρ*_r_, of a cell equilibrated to a hypotonic solution and subsequently returned to isotonic conditions is given by the function,

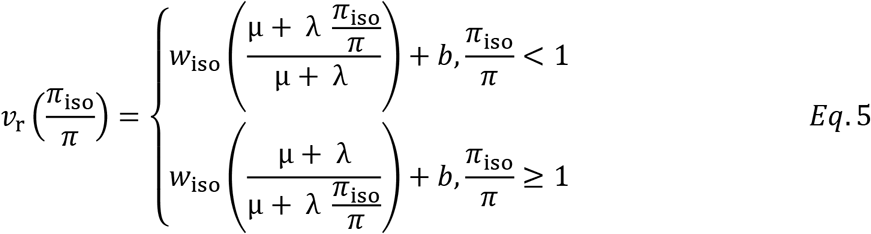

which assumes no leakage during return to isotonic conditions after swelling. However, during return from hypertonic conditions this model does predict leakage and therefore is presented as a piecewise function.

Intracellular solute control is an effective method of regulated osmotic challenge but has associated physiological challenges. The change in cell membrane permeability may result in alteration of internal ion concentration balances [18,78], the loss of carbohydrates [84], and the total arrest of metabolic activity [85]. A method of mitigating volume change while forgoing these physiological challenges may be found in mechanically resisting deformation.

#### 4.1.3 Turgor Model

Mechanical resistance to hypoosmotic challenge is a common strategy through the Plantae, Fungi, Protista, Archaea and Bacteria kingdoms, and turgor models of the elastic resistive pressure is traditionally included in the hydrostatic pressure difference for the water flux model [86–88]. The flux of water across the cell membrane may be written in terms of chemical potential, Δ*Ψ*, which in turn can be written as the difference in both hydrostatic pressure, Δ*P*, and osmotic pressure Δ*Π*, across the membrane, written as Δ*Ψ* = Δ*P* – Δ*Π*. During swelling in hypotonic conditions for plants and yeast cells, strain in the cell wall is proportional to hydrostatic pressure at the cell membrane such that as the strain increases, the hydrostatic pressure increases until Δ*P* = Δ*Π* (reducing the net flux of water into the cell) [86,89]. Mechanical resistance to volumetric change for animals is generally thought to be minimal since lipids in the cell membrane have relatively little resistive force [4,18,90,91]. However, the lipid membrane is directly incorporated into the cytoskeletal cortex which increases the mechanical stability and resistance to deformation of the cell, collectively called the membrane-cortex [90,92,93].

Indeed, analogous to the membrane-cortex structure is the nuclear lamina. The nuclear lamina stabilizes the nucleus during osmotic shock during hypotonic challenge [38,94]. Experimentally, in the case of the nuclear lamina, nuclear volume appears to behave linearly in the hypertonic region and seems not to noticeably resist shrinkage, though it does elastically resist swelling [38]. Similar to the nuclear lamina, the cytoskeleton cortex stabilizes the lipid membrane and provides resistance to deformation [90,92,93]. Making the argument that the membrane-cortex behaves similarly to the nuclear lamina, it is then reasonable to assume linear shrinkage (or insignificant resistance to shrinkage) while asserting an elastic resistance to swelling [31,38,95].

Cells and tissues may have swelling limited by an extracellular matrix as well. For example, mammalian oocytes and embryos are surrounded by the zona pellucida, a glycoprotein shell, shown to resist deformation [72]. The yeast cell wall has been shown to have a Young’s modulus of 20.3 MPa [86], whereas bovine zona pellucida have Young’s modulus reported ranging from 1 kPa to 600 kPa [71,72,96]. Mammalian oocytes may have a smaller yet still significant resistance to swelling when compared to yeast cells, at least when the oocyte swells to the enclosed volume of the zona pellucida. In fact, this resistance may describe the observed nonlinear behaviour of some BvH plots such as golden hamster oocytes [52].

Complex models of turgor exist, however, these require a number of parameters determined from experiments beyond those typical of BvH plots [4,6,8,88,94]. Therefore, here we develop a simple short-term turgor model, using two key assumptions: the first is that the cell behaves linearly as with respect to the BvH relation during hypertonic challenge (i.e. the cell does not elastically resist shrinkage), and second, while swelling past the “turgor point”, the cell elastically resists deformation as a thin shell. The “turgor point” is defined as the volume of the cell at which the turgor pressure of the cell is equal to zero, i.e. there is no elastic resistance, but if the cell increases in volume the elastic resistance increases. Note that more details of the derivation of this model can be found in the appendix. Because few mammalian cell BvH datasets we found had more than 5 points, we chose to set to the turgor point to be the isotonic point to avoid an extra parameter and overfitting. Moreover, we do not consider thick-shelled deformation, nor the possible hydro-gel-like properties of cell deformation [38]. The Poisson ratio *ν* of membranes may be assumed to be 0.5 [97,98]. Assuming an isotropic and homomorphic shell at the cell boundary, and linear elastic deformation with respect to the radius, a classical thin shell elastic one-dimensional equation may be used. The strain in the shell is then proportional to change in pressure and thus change in hydrostatic pressure, Δ*P*, at the cell membrane. We note the thin and thick shelled elastic problem is common in the field of elasticity theory and found covered in most textbooks [99]. The piecewise turgor function takes the form,

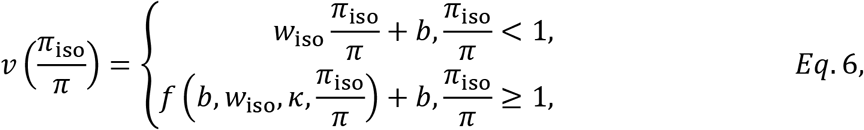

where the function *f* returns the real positive solution for *w* with respect to the implicit equation,

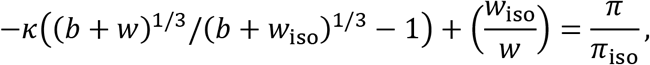

Where *κ* is a lumped elasticity term such that *κ* = 2 *K τ*/ (1 – *ν*) *r*_i_ *R T π*_iso_, and *K* is the Young’s modulus, *τ* is the combined thickness of the cell membrane and cortex [4], and *r*_i_ is the reference radius of the cell. This piecewise model (*Eq*. 6) reduces to a linear model (*Eq*. 2) when *κ* = 0, and is nonlinear in hypotonic conditions when *κ* > 0. The three constants, *b*, *w*_iso_, and *κ* can be fitted simultaneously to a BvH plot. Furthermore, this model makes the assumption *π*_iso_/*π* ≅ *c*_iso_/*c* in the hypotonic region and thus may be used with both osmolality and osmolarity. Finally, while more complex mechanical models exist, *Eq*. 6 provides a robust approximation using a single parameter that may be fitted solely from a BvH plot. See the appendix for the development of *Eq*. 6.

This turgor model differs from the leak model in that it assumes all volume regulation is mechanical and hence there should be no change to the quantity of intracellular solutes. This means that after exposure to anisosmotic conditions the cell will return to the original cell volume in isotonic conditions. To our knowledge, there are no Boyle van ’t Hoff studies in the literature that also report on “return to isotonic” volume responses, so a differentiation between the two models with no further experimentation can only be asserted by “goodness of fit” to the data and lack of autocorrelation.

#### 4.1.4 Turgor-Leak Model

Both the turgor model and the leak model assume a single mechanism of osmoregulation (mechanical resistance or ion leakage). On the contrary, Hua and colleagues [42] suggest a tiered combination of mechanical resistance and osmolytes leakage for cells to mitigate osmotic challenges. The cytoskeleton is theorized to retain most of the tension generated by an osmotic challenge [42,60,91]. The cytoskeleton is both mechanically protective and a mechanical messenger for the cell [92,100,101]. When the cytoskeleton starts to break down, local cortical and membrane tension rises and mechanosensitive channels embedded in the lipid membrane and/or directly connected to the cytoskeletal cortex are activated [42]. Expulsion of internal osmolytes is a way to re-establish equilibrium without an over expansion of cell volume, protecting from lysis. However, the loss of internal osmolytes may be energetically expensive for the cell to reobtain, potentially leading to cell death. It has been shown the cells may fully suspend metabolism while losing a majority of ATP as well as other complex molecules such as sugars [84,85]. Furthermore, without the cytoskeleton, cell survival in hypotonic challenge is dramatically impaired, where 84% of monolayer MDCK cells may survive very hypotonic exposure (~3 mOsm/kg), treatment with cytochalasin D (which denatures the cytoskeleton) results in no surviving cells after exposure [85]. It seems plausible that the cell initially resists osmotic challenges via mechanical resistance then becomes increasingly permeable to ions followed by organic osmolytes. A combination of ion channels, gates, pumps, and a dynamic cytoskeleton may provide improved predictive power modeling cellular response to osmotic challenge [4].

Indeed, Jiang & Sun [4] present a model of mechanical resistance and ion leakage with a total of 11 parameters (while exluding ion specific channels/gates and membrane regulation). Here we present a simplified mechanical resistance and ion-osmolyte leak model for short time scales such that ion pumps may be safely removed (since flux of ions via ion channels is a few order of magnitudes faster than ion pumps). Furthermore, we assume while the membrane-cortex is under tension all ions and osmolytes are free to flow through the mechanosenstive channels such that the flux of ions is proportional to the chemical potential gradient of ions across the membrane which is propotional to the molality gradient Δ*m* in hypotonic conditions. For simplicity all ions and osmolytes are grouped together to form the total molality gradient. Since we are grouping all ions and osmolytes together then *m*_e_ ≈ *π*_e_. Furthermore, since the flux of water across the membrane is proportional to the chemical potential gradient, which itself is proportional to the difference of hydrostatic gradient Δ*P* and osmotic pressure gradient Δ*Π*, then the simplified turgor-leak model takes the form,

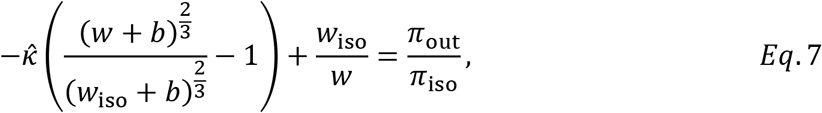

Where *A* is the surface area of the cell, *L*_p_ is hydraulic conductivity across the cell membrane, *P_n_* is the lumped membrane permeability (including of both channels and gates), and other parameters are as defined above.

Interestingly, when combining both hydrostatic resistance and ion leakage, the cell volume will return to its rest state passively over time (without the necessity of active pumps). This is because when *dn*/*dt* = 0, then *π*_e_ = *n*/(*ρW*), meaning *RT*[*π*_e_ – *n*/(*ρW*)]= 0 and thus water flux is driven by hydrostatic pressure such that

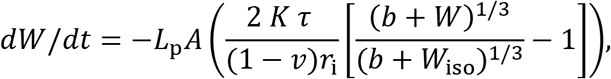

Forcing water out of the cell when the cell is swollen (*W* > *W*_iso_). Since water is forced out of the cell, then *π*_e_ < *n*/(*ρW*), and thus ions/osmolytes leave the cell to restore equilibrium. This continues until the resting equilbrium occurs such that *π*_e_ = *n*/(*ρW*) and *W* = *w*_iso_. Note that *w*_iso_ is in reference to the “resting” point of the cell with respect to mechanical resistance, and in our case for simplicity is set to the isotonic point. A dynamic resting point using a plastic and self regulating membrane-cortex may improve the predicted equilibrium volume at the expense of a more complex model [18,60]. Notably, this passive reduction in volume does not occur with the leak model or turgor model by themselves since a stable equilibrium is reached for both models. Interestingly, the same argument of a passive volume regulation (without the need for active pumps to return to the resting volume) may be used during hypertonic shrinkage if the cell mechanically resists shrinkage. If high membrane tensions form membrane-cortex compression or high curvature in the lipid membrane results in increased ion-osmolyte permeability, then the cell may use the combination of mechanical resistance and leakage to return to initial volume without the use of active pumps. The main difference is that the hydrostatic pressure is in the opposite direction, meaning water is forced into the cell followed by an influx of ions/osmolytes to restore equilibrium conditions. The resting point then is reached when *π*_e_ = *n*/(*ρW*) and *W* = *W*_iso_. Finally, *Eq*. 7 predicts both volume of the cell throughout time but also return to isotonic volume after exposure to anisotonic conditions. Using both volume during osmotic challenge as well as return to isotonic conditons provides a methodology to test ion leakage as well as mehanical resistance simultaniously. Therefore, *Eq*. 7 may be fit to a BvH plot in combination with a return to isotonic conditions plot. Notably, *Eq*. 7 predicts cells in hypertonic conditions to have larger return volume than initial volume, which is in direct contrast to the leak model which predicts a smaller return volume.

### 4.2 Meta Analysis Data Collection

During the summer of 2019, we collected volume versus inverse osmolality plots by searching the phrases “boyle van’t hoff”, “boyle vant hoff’ and “Ponder’s Plot” in Google Scholar, with and without quotation marks. We first examined the title of all the articles and selected all articles that were remotely likely to contain BvH data. Then we combed through these articles and selected articles that had original BvH data. Finally, we thoroughly reviewed each paper and selected papers with “normal BvH data”. We define “normal BvH data” to be data of cell volume equilibrated in a solution of relatively nonpermeating solutes as a function of osmolality, osmolarity, or osmotic pressure. Further, data were restricted to experiments conducted at room temperature or physiological temperature without any chemical alterations (e.g. the addition of Lactruculin D). We did not collect from articles that stated equilibrium was not reached.

Mean data points were extracted from figures with the use of webPlotDigitizer (https://automeris.io/WebPlotDigitizer/). Data were normalized with respect to stated values in the respective articles. Note that the extracted mean data does not include variance for the respective data points. We identified 137 potential articles ranging from publication years of 1964 to 2019. We composed three groups of datasets. **Group 1** had 6 or more data points for each dataset and an inverse relative osmolality range of 0.5 to 2 or more, resulting in a total of 28 datasets containing 247 mean data points. Note that for cells with an isotonic point of 290 mOsm/kg, this is equivalent to assuring a range of at least 145 mOsm/kg to 580 mOsm/kg.

Supplementary Material 1 provides more information regarding these datasets. **Group 2** had 6 or more data points for each dataset for the hypertonic region (relative inverse osmolality ranging from 0 to 1) resulting in a total of 22 datasets and 167 data points. Supplementary Material 2 provides more information regarding these datasets. **Group 3** only had the restriction range of 0.5 to 2 relative inverse osmolality or more resulting in a total of 44 datasets and 326 data points. Supplementary Material 3 provides more information regarding these datasets. Our total collection spanned 14 animal types and 26 cell types with 38 unique animal-cell type combinations (e.g. “Human hepatocyte”, or “Bovine oocyte”). All meta-analysis data are presented in terms of relative inverse osmotic pressure with the assumption that, *Π*_iso_/*Π* ≅ *π*_iso_/*π* ≅ *c*_iso_/*c*, which is a necessary step for comparing trends between fitted datasets.

### 4.3 Linear fitting Autocorrelation Analysis

A method of determining a response is linear is analyzing the autocorrelation of the residuals of a linear model [102–104]. Thus the Durbin Watson (DW) score was used to measure lag one autocorrelation of each model against volume-osmolality data and ranges from 0 to 4 [102,104]. Note that a DW score close to 2 is considered not autocorrelated while values close to 0 and 4 are positively and negatively autocorrelated respectively. Since negative autocorrelation are a signal of spatial-temporal oscillation, and since our preliminary observations indicated a large majority of nonlinear BvH plots had positive autocorrelation, we focused on testing the hypothesis of positive autocorrelation for this study.

Each dataset was independently fit with the nondimensional BvH relation (*Eq*. 2) through linear regression, using Mathematica’s “LinearModelFit” function [105]. Savin & White (1977) provide critical bounds for datasets of 6 or more data points while datasets of 5 or less are ignored due to high variability and low power [103]. Using the critical lower bounds derived from Savin & White (1977), we counted the number of datasets that were below the critical bound (using α=0.05) for Group 1 (hyper- and hypotonic data) and Group 2 (hypertonic data). We then conducted an exact binomial test to determine if the observed number of rejected null hypotheses is significantly larger than expected (5%) for each group.

### 4.4 Nonlinear fitting and comparison

All nonlinear models were fit to each respective dataset using the “NonLinearModelFit” function in Mathematica [105]. We used three methods of comparison between models for Group 3 datasets. First, we compared total mean squared error (TMSE). This was be done by calculating the mean squared error (MSE) for each dataset for a given model and then summing each MSE together to provide the total mean squared error for the respective model across Group 3.

Second, we compared mean adjusted R^2^ values across nonlinear models by calculating the associated adjusted R^2^ for each dataset given for each model. We conducted a Mann-Whitney U to determine if there was a difference of median adjusted R^2^ between nonlinear models. Finally, we counted the ranked AICc score for each model. This was be done by taking each dataset and ranking each model in terms of the lowest AICc, with the lowest value being ranked 1. The total count of each model being ranked 1 across datasets were presented (i.e. the number of times a given model has the lowest AICc). Note we used AICc and not AIC or BIC as many datasets have low number of data points. Furthermore, note that if the number of data points is too low, with respect to the number of fitted parameters, then the AICc value is given a score of positive infinity (ruling it out as a competing model for the respective dataset).

### 4.5 Cell Culture

The HepG2 hepatocellular carcinoma cell line was purchased from the American Tissue Culture Collection (ATCC; HepG2/C3A, Catalog no. CRL-10741). HepG2 cells were cultured in Eagle’s Minimum Essential Medium (MEM) with 1.5 g/L sodium bicarbonate, non-essential amino acids, L-glutamine, and sodium pyruvate (Sigma-Aldrich, Canada), supplemented with 10% FBS at 37 °C and 5% CO_2_ atmosphere. Media was changed every 48 h. When they reached 60-70% confluency, cells were harvested with 0.25% Trypsin-EDTA solution, centrifuged at 300 g for 5 minutes and suspended at a density of 1x 10^6^ cells/ml in anisotonic solutions.

### 4.6 Test Solutions

Hyper- and hypotonic solutions containing only non-permeating solutes were prepared by diluting phosphate-buffered saline (PBS; Gibco-Life Technologies) with distilled water to create solution osmolalities of 80, 100, 200, 400, 600 mOsm/kg and isotonic (290 mOsm/kg).

Osmolalities were measured by freezing point depression osmometer (Fiske Associates, Norwood, Massachusetts, USA).

### 4.7 Measurement of Cell Volume During Osmotic Runs

Before equilibration with anisotonic solutions, trypsinized HepG2 cells were resuspended in fresh culture medium and equilibrium runs were carried out at room temperature (21 °C), where cells were injected into hypo and hypertonic solutions of 80, 100, 200, 400, 600 mOsm/kg. Cells were left to equilibrate for 6 minutes (an experimental duration time estimated from preliminary experiments examining the evolution of the cell volume distribution with time). Using optical microscopy, photomicrographs were imaged with a 20X/long working distance objective NA 0.4. After equilibration with anisotonic solutions, cells were restored to isotonic condition to determine their return to isotonic volume. The change in the shape and size of hepatocytes were registered photographically and the diameter of the cells were extracted via ImageJ. Cell volumes were estimated by using formula of the sphere while trypan blue was used to exclude the dead cells. Cell volumes were normalized with respect to isotonic cell volume of 1769±58.6 μm^3^ (n=100).

## Supporting information

Supplemental Material 3 (Group 3 Data)

Supplemental Material 2 (Group 2 Data)

Supplemental Material 1 (Group 1 Data)

## 5 Appendix Derivation of the Turgor Model

Water flux across membrane is proportional to chemical potential gradient, ΔΨ. The chemical potential on the outside of the cell can be written as *Ψ*_out_ = *P*_out_ – *Π*_out_, where *P*_out_ is the extracellular hydrostatic pressure, and *Π*_out_ is the extracellular osmotic pressure. The chemical potential inside the cell can be written as *Ψ*_in_ = *P*_in_ – *Π*_in_, such that the chemical potential gradient across the cell membrane is Δ*Ψ* = Δ*P* – Δ*Π*, where Δ*P* = *P*_in_ – *P*_out_, and Δ*Π* = *Π*_in_ – *Π*_out_. The flux of water, *dw*/*dt*, across the cell membrane any moment is

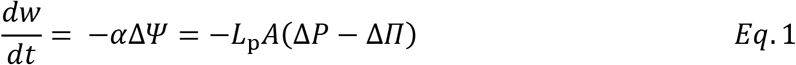

Where *α* is a rate constant equal to the hydraulic conductivity *L*_p_, and the surface area of the cell *A*.

Hydrostatic pressure is in mechanical equilibrium with the mechanical stress of the cell boundary matrix (either cell wall, zona pellucida, or membrane-cortex). The stress may be modelled by general viscoelastic models [4], polymer mechanics [88], and elastic cortex models [38,106]. However, the simple solution of a thin isotropic linear elastic model provides a robust estimate of mechanobiological behaviours in the context of both the zona pellucida and the membrane-cortex for the respective cells. The radial strain *ε*, is related to stress in a thin-walled sphere by,

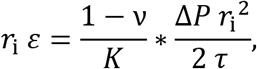

where pressure differential Δ*P* across the membrane is equal to hydrostatic pressure differential, *r*_i_ is the reference radius of the cell, *K* is the Young’s modulus, *ν* is the poisson ratio (taking the value of 0.5), *τ* is the thickness of the thin-wall structure (e.g. the membrane-cortex, the cell wall, or the zona pellucida). Letting normal strain be defined as the change in length, then *ε* = (*r* – *r*_i_)/*r*_i_, when *r* is the new radius of the cell. We can describe changes to the hydrostatic pressure at the cell boundary by writing pressure in terms of the above equation terms of radial change,

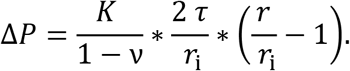

Letting total cell volume *V* be composed of intracellular water volume *W* and non-osmotic volume *V*_b_, the above equation can be written in terms of relative volume, letting the reference radius *r*_i_ = *r*_iso_, the normalized equation takes the form,

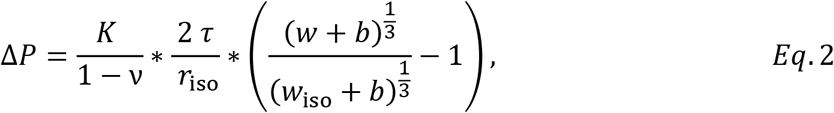

where the non dimensional volume fractions are defined by *w*_iso_ = *W*_iso_/*V*_iso_, *w* = *W*/*V*_iso_, and *b* = *V*_b_/*V*_iso_.

Noting that Δ*Π* = *Π*_in_ – *Π*_out_ = *RT*(*π*_in_ – *π*_out_) = *RT*(*n*/*ρW* – *π*_out_), and *n* = *ρW π*_iso_, then normalizing with respect to *V*_iso_ yields,

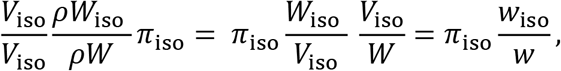

thus,

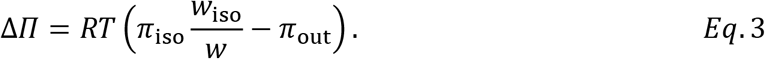

Substituting *Eq*. 2 and *Eq*. 3 into *Eq*. 1 yields the equation,

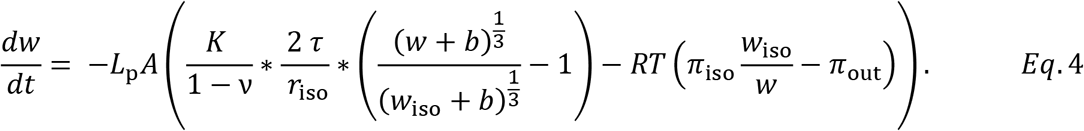

At equilibrium, the net flux of water across the membrane is zero, thus *dw*/*dt* = 0, and thus the solution of *Eq*. 4 at equilibrium is,

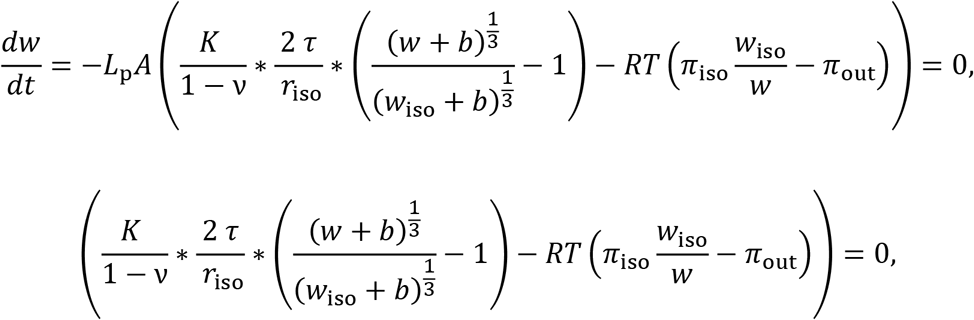

and simplifying,

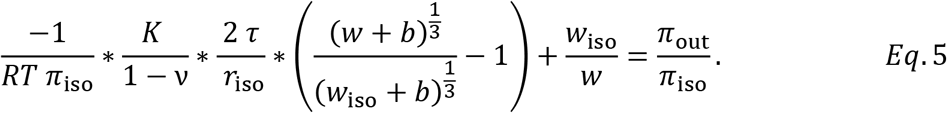

Letting 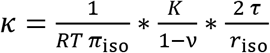, we arrive at the normalized implicit solution of cell water volume at equilibrium as

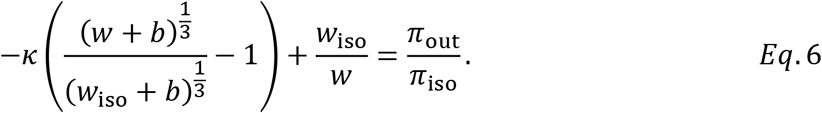

The explicit solution for *w* in *Eq*. 6 may be determined via a numerical root finding method given *w*_iso_, *b*, and *κ* as fitting parameters. The total cell volume then is the water volume derived from *Eq*. 6 plus the non-osmotic volume of the cell.

It is worth noting that another method of modelling elastic cortex is with respect to surface area of the cell [4]. Hydrostatic pressure gradient is written in terms of deformed surface area as,

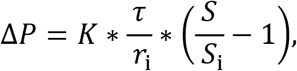

Where *K* is a lumped elastic modulus containing the Young’s modulus and the Poisson ration.

The above equation may be written in terms of normalized volume as

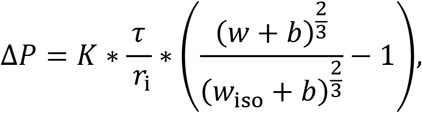

with a resulting implicit equation at equilibrium as

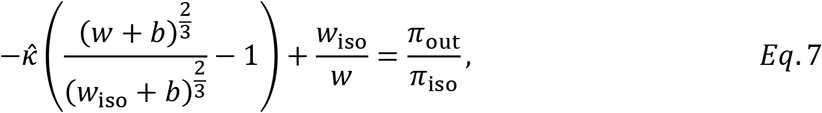

Where 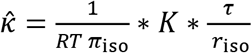. Plotting *Eq*. 6 or *Eq*. 7 in a BvH plot results in almost entirely overlapping fitted models, but with 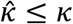. Assuming a membrane-cortex thickness of 0.5 μm, asserting a Poisson ratio of 0.5 for both models, and fitting both models to the HepG2 BvH plot in this paper, then Eq. 6 has a fitted Young’s modulus of 6.265 MPa while *Eq*. 7 has a fitted a fitted Young’s modulus of 5.817 MPa, with an absolute error of 7%.

While asserting no shrinkage resistance simplifies the model (not requiring a fitted shrinkage elastic parameter), fitting both shrinking and swelling by a single elastic parameter works well. However, this model is still outperformed by assuming only swelling resistance, especially for datasets with a low number of data points. While the turgor model presented in this paper is shown to be equivalent to the mechanical resistance model presented by Jiang and Sun [4], linearly elastic models assume relatively small strains, which might not necessarily be the case when the cell swells to twice its volume (an increase of 26% for the radius). Using polymer elastic principles [88], and by experimentally identifying when the oocyte membrane presses uniformly against the zona pellucida for identification of the turgor point, may provide an improved turgor model. However, we point out the wide application of the current simple thin shell model fitting well to many cell types (Figure 9). Although not covered in this paper, our turgor model may also be used for plant [88,89], yeast [86], mitochondria [31,95], nucleus [38], and bacterial [107] cells with robust fitting to the associated BvH plots. Further work may seek to determine and experimentally compare between competing mechanical resistance models.

## Notes

### Competing Interest Statement

The authors have declared no competing interest.

