## Supplemental Material 3 (Group 3 Data) for "Meta-analysis and experimental re-evaluation of the Boyle van ‘t Hoff relation with osmoregulation modelled by linear elastic principles and ion-osmolyte leakage"

| Table ID | Model | b | wiso | nonlinear Const. | MSE | adj$R^2$ | AICc | Animal Type | Cell Type | # data points | Min & Max Range | Journal Article |
|---|---|---|---|---|---|---|---|---|---|---|---|---|
| 1,1 | BvH (Hyper) | 0.154 | 0.851 | | | | | Golden Hamster | Pancreatic islet cell | 6 | 0.15<br>2.01 | Benson et al.<br>1993.<br>Cell Transplantation |
| | BvH | 0.222 | 0.702 | | $2.501 \times 10^{-3}$ | 0.98571 | -0.48604 | | | | | |
| | Leak | 0.149 | 0.857 | $1.19 \times 10^{-3}$ | $1.233 \times 10^{-5}$ | 0.99997 | -1.6339 | | | | | |
| | Turgor | 0.155 | 0.848 | $6.52 \times 10^{-1}$ | $1.459 \times 10^{-5}$ | 0.99997 | -0.62393 | | | | | |
| 1,2 | BvH (Hyper) | 0.716 | 0.297 | | | | | Bovine | Spermatozoa | 7 | 0.502<br>3. | Basse et al.<br>2012.<br>Theriogenology |
| | BvH | 0.731 | 0.262 | | $1.494 \times 10^{-3}$ | 0.96058 | -11.326 | | | | | |
| | Leak | 0.73 | 0.263 | $2.94 \times 10^{-8}$ | $1.494 \times 10^{-3}$ | 0.99795 | 3.2365 | | | | | |
| | Turgor | 0.731 | 0.262 | $2.18 \times 10^{-5}$ | $1.494 \times 10^{-3}$ | 0.99795 | 3.2365 | | | | | |
| 1,3 | BvH (Hyper) | 0.265 | 0.705 | | | | | Human | Chondrocyte | 11 | 0.542<br>9.53 | Bush and Hall<br>2005.<br>Journal of cellular physiology |
| | BvH | 0.842 | 0.0899 | | $4.144 \times 10^{-2}$ | 0.50257 | 5.8335 | | | | | |
| | Leak | 0.151 | 0.874 | $8.07 \times 10^{-3}$ | $2.408 \times 10^{-3}$ | 0.9971 | -19.931 | | | | | |
| | Turgor | 0.189 | 0.82 | 3.16 | $1.953 \times 10^{-3}$ | 0.99765 | -22.235 | | | | | |
| 1,4 | BvH (Hyper) | 0.219 | 0.776 | | | | | Lobster | axon | 8 | 0.32<br>4. | Freeman et al.<br>1966.<br>The Journal of general physiology |
| | BvH | 0.54 | 0.334 | | $3.377 \times 10^{-2}$ | 0.7845 | 7.8982 | | | | | |
| | Leak | 0.146 | 0.919 | $1.1 \times 10^{-2}$ | $2.405 \times 10^{-3}$ | 0.99655 | -3.4437 | | | | | |
| | Turgor | 0.172 | 0.879 | 1.87 | $1.785 \times 10^{-3}$ | 0.99744 | -5.8319 | | | | | |
| 2,1 | BvH (Hyper) | 0.298 | 0.715 | | | | | Human | hematopoietic progenitor cells | 8 | 0.215<br>1.99 | Gao et al.<br>1998.<br>Cryobiology |
| | BvH | 0.21 | 0.87 | | $8.947 \times 10^{-3}$ | 0.96167 | -2.7269 | | | | | |
| | Leak | 0.206 | 0.874 | $1.91 \times 10^{-9}$ | $8.947 \times 10^{-3}$ | 0.98805 | 7.065 | | | | | |
| | Turgor | 0.21 | 0.87 | $9.19 \times 10^{-7}$ | $8.947 \times 10^{-3}$ | 0.98805 | 7.065 | | | | | |
| 2,2 | BvH (Hyper) | 0.493 | 0.512 | | | | | Human | Spermatozoa | 4 | 0.316<br>1.97 | Gilmore et al.<br>1995.<br>Biology of reproduction |
| | BvH | 0.515 | 0.472 | | $1.587 \times 10^{-4}$ | 0.99745 | ∞ | | | | | |
| | Leak | 0.491 | 0.514 | $2.95 \times 10^{-4}$ | $3.562 \times 10^{-5}$ | 0.99986 | ∞ | | | | | |
| | Turgor | 0.493 | 0.512 | $2.78 \times 10^{-1}$ | $3.56 \times 10^{-5}$ | 0.99986 | ∞ | | | | | |
| 2,3 | BvH (Hyper) | 0.811 | 0.195 | | | | | Equine | Spermatozoa | 5 | 0.333<br>4. | Gonzalez-Fernandez et al.<br>2012.<br>Theropgenology |
| | BvH | 0.778 | 0.289 | | $3.77 \times 10^{-3}$ | 0.96778 | 16.841 | | | | | |
| | Leak | 0.72 | 0.371 | $3.56 \times 10^{-4}$ | $2.674 \times 10^{-3}$ | 0.996 | ∞ | | | | | |
| | Turgor | 0.72 | 0.368 | $2.93 \times 10^{-1}$ | $2.287 \times 10^{-3}$ | 0.99658 | ∞ | | | | | |
| 2,4 | BvH (Hyper) | 0.303 | 0.812 | | | | | Human | Chondrocyte | 9 | 0.537<br>3. | Guilak<br>2000.<br>Bioreheology |
| | BvH | 0.257 | 0.891 | | $4.36 \times 10^{-3}$ | 0.99054 | -12.314 | | | | | |
| | Leak | 0.181 | 0.992 | $2.2 \times 10^{-4}$ | $3.757 \times 10^{-3}$ | 0.99762 | -6.0661 | | | | | |
| | Turgor | 0.193 | 0.975 | $8.26 \times 10^{-2}$ | $3.627 \times 10^{-3}$ | 0.9977 | -6.3841 | | | | | |
| 3,1 | BvH (Hyper) | 0.292 | 0.699 | | | | | Human | CD34+ cell | 10 | 0.161<br>1.95 | Hunt et al.<br>2003.<br>Cryobiology |
| | BvH | 0.273 | 0.731 | | $1.581 \times 10^{-3}$ | 0.99165 | -25.884 | | | | | |
| | Leak | 0.27 | 0.734 | $5. \times 10^{-9}$ | $1.581 \times 10^{-3}$ | 0.9978 | -19.549 | | | | | |
| | Turgor | 0.273 | 0.731 | $2.81 \times 10^{-6}$ | $1.581 \times 10^{-3}$ | 0.9978 | -19.549 | | | | | |
| 3,2 | BvH (Hyper) | 0.102 | 0.926 | | | | | Dog | notochordal cell | 8 | 0.335<br>7.81 | Hunter et al.<br>2007.<br>Mol Cell Biomech. |
| | BvH | 0.603 | 0.351 | | $3.143 \times 10^{-2}$ | 0.94688 | 7.3248 | | | | | |
| | Leak | 0.307 | 0.633 | $5.28 \times 10^{-4}$ | $1.032 \times 10^{-2}$ | 0.99398 | 8.2105 | | | | | |
| | Turgor | 0.372 | 0.551 | $1.35 \times 10^{-1}$ | $1.31 \times 10^{-2}$ | 0.99236 | 10.116 | | | | | |
| 3,3 | BvH (Hyper) | 0.302 | 0.698 | | | | | Crab | Muscle Fiber | 10 | 0.33<br>2.07 | Lang and Gainer<br>1969.<br>The Journal of General Physiology |
| | BvH | 0.382 | 0.588 | | $2.44 \times 10^{-3}$ | 0.97504 | -21.547 | | | | | |
| | Leak | 0.244 | 0.792 | $4.2 \times 10^{-3}$ | $7.422 \times 10^{-4}$ | 0.99912 | -27.113 | | | | | |
| | Turgor | 0.262 | 0.767 | $8.83 \times 10^{-1}$ | $6.115 \times 10^{-4}$ | 0.99927 | -29.05 | | | | | |
| 3,4 | BvH (Hyper) | 0.253 | 0.753 | | | | | Human | Lymphocyte | 17 | 0.103<br>2.03 | Law et al.<br>1983.<br>Cryobiology |
| | BvH | 0.244 | 0.779 | | $2.165 \times 10^{-3}$ | 0.98492 | -48.086 | | | | | |
| | Leak | 0.236 | 0.794 | $7.3 \times 10^{-5}$ | $2.153 \times 10^{-3}$ | 0.99603 | -44.52 | | | | | |
| | Turgor | 0.237 | 0.796 | $6.29 \times 10^{-2}$ | $2.134 \times 10^{-3}$ | 0.99606 | -44.67 | | | | | |
| 4,1 | BvH (Hyper) | 0.407 | 0.607 | | | | | Hamster | Pancreatic islet cell | 6 | 0.201<br>3.35 | Lui et al.<br>1995.<br>Cryobiology |
| | BvH | 0.621 | 0.193 | | $1.567 \times 10^{-2}$ | 0.68523 | 10.526 | | | | | |
| | Leak | 0.4 | 0.62 | $1.29 \times 10^{-2}$ | $3.925 \times 10^{-4}$ | 0.99902 | 19.128 | | | | | |
| | Turgor | 0.404 | 0.614 | $1.16 \times 10^{1}$ | $3.7 \times 10^{-4}$ | 0.99908 | 18.774 | | | | | |
| 4,2 | BvH (Hyper) | 0.2 | 0.955 | | | | | Mouse | Blastocyte (with zona) | 7 | 0.174<br>4.93 | Mazur and Schneider<br>1985.<br>Cell Biophysics |
| | BvH | 0.446 | 0.656 | | $4.82 \times 10^{-2}$ | 0.95177 | 12.993 | | | | | |
| | Leak | 0.155 | 1.03 | $7.23 \times 10^{-4}$ | $1.502 \times 10^{-2}$ | 0.99461 | 19.394 | | | | | |
| | Turgor | 0.181 | 0.966 | $2.1 \times 10^{-1}$ | $1.295 \times 10^{-2}$ | 0.99535 | 18.358 | | | | | |
| 4,3 | BvH (Hyper) | 0.488 | 0.331 | | | | | Mouse | Embryo eight cell (with zona) | 5 | 0.151<br>4.9 | Mazur and Schneider<br>1985.<br>Cell Biophysics |
| | BvH | 0.563 | 0.384 | | $4.793 \times 10^{-2}$ | 0.87781 | 29.554 | | | | | |
| | Leak | 0.32 | 0.916 | $2.22 \times 10^{-3}$ | $5.797 \times 10^{-3}$ | 0.99227 | ∞ | | | | | |
| | Turgor | 0.341 | 0.851 | $9.4 \times 10^{-1}$ | $4.653 \times 10^{-3}$ | 0.99379 | ∞ | | | | | |
| 4,4 | BvH (Hyper) | 0.171 | 0.827 | | | | | Bovine | Morulae | 10 | 0.118<br>2.86 | Mazur and Schneider<br>1985.<br>Cell Biophysics |
| | BvH | 0.134 | 0.853 | | $4.415 \times 10^{-3}$ | 0.99009 | -15.618 | | | | | |
| | Leak | 0.13 | 0.857 | $1.34 \times 10^{-9}$ | $4.415 \times 10^{-3}$ | 0.99695 | -9.2829 | | | | | |
| | Turgor | 0.134 | 0.853 | $4. \times 10^{-7}$ | $4.415 \times 10^{-3}$ | 0.99695 | -9.2829 | | | | | |
| 5,1 | BvH (Hyper) | 0.219 | 0.782 | | | | | Mouse | Ova | 13 | 0.0846<br>5.6 | Oda et al.<br>1992.<br>Journal of reproduction and fertility |
| | BvH | 0.387 | 0.514 | | $4.065 \times 10^{-2}$ | 0.92998 | 4.0956 | | | | | |
| | Leak | 0.189 | 0.876 | $7.56 \times 10^{-4}$ | $3.849 \times 10^{-3}$ | 0.99692 | -21.975 | | | | | |
| | Turgor | 0.202 | 0.838 | $3.07 \times 10^{-1}$ | $2.612 \times 10^{-3}$ | 0.99791 | -27.015 | | | | | |
| 5,2 | BvH (Hyper) | 0.775 | 0.248 | | | | | Horse | Spermatozoa | 7 | 0.498<br>2. | Oldenhof et al.<br>2011.<br>Theriogenology |
| | BvH | 0.467 | 0.682 | | $7.127 \times 10^{-3}$ | 0.93117 | -0.38623 | | | | | |
| | Leak | 0.465 | 0.685 | $5.43 \times 10^{-9}$ | $7.127 \times 10^{-3}$ | 0.99315 | 14.176 | | | | | |
| | Turgor | 0.467 | 0.682 | $4.25 \times 10^{-6}$ | $7.127 \times 10^{-3}$ | 0.99315 | 14.176 | | | | | |
| 5,3 | BvH (Hyper) | 0.219 | 0.852 | | | | | Hamster | Oocyte (with zona) | 9 | 0.0471<br>5.99 | Oungoulian et al.<br>2011.<br>Proceedings of the ASME 2011 Summer Bioengineering Conference |
| | BvH | 0.62 | 0.174 | | $4.829 \times 10^{-2}$ | 0.64205 | 9.3285 | | | | | |
| | Leak | 0.234 | 0.796 | $6.02 \times 10^{-3}$ | $2.344 \times 10^{-3}$ | 0.99654 | -10.313 | | | | | |
| | Turgor | 0.238 | 0.788 | 4.44 | $2.724 \times 10^{-3}$ | 0.99598 | -8.962 | | | | | |
| 5,4 | BvH (Hyper) | 0.249 | 0.754 | | | | | Mouse | cortex neuron | 7 | 0.23<br>2.31 | Paynter et al.<br>2009.<br>Cryobiology |
| | BvH | 0.311 | 0.651 | | $7.266 \times 10^{-3}$ | 0.9614 | -0.25165 | | | | | |
| | Leak | 0.205 | 0.865 | $1.24 \times 10^{-3}$ | $2.559 \times 10^{-3}$ | 0.99569 | 7.006 | | | | | |
| | Turgor | 0.21 | 0.855 | $7.53 \times 10^{-1}$ | $2.095 \times 10^{-3}$ | 0.99647 | 5.6063 | | | | | |
| 6,1 | BvH (Hyper) | 0.253 | 0.726 | | | | | Mouse | cortex neuron | 7 | 0.232<br>2.32 | Paynter et al.<br>2009.<br>Cryobiology |
| | BvH | 0.26 | 0.725 | | $1.595 \times 10^{-3}$ | 0.99301 | -10.866 | | | | | |
| | Leak | 0.223 | 0.797 | $3.04 \times 10^{-4}$ | $1.077 \times 10^{-3}$ | 0.99832 | 0.95014 | | | | | |
| | Turgor | 0.226 | 0.794 | $1.76 \times 10^{-1}$ | $9.531 \times 10^{-4}$ | 0.99851 | 0.091804 | | | | | |
| 6,2 | BvH (Hyper) | 0.237 | 0.771 | | | | | Mouse | ventral mesencephalon nerual cell | 7 | 0.232<br>2.32 | Paynter et al.<br>2009.<br>Cryobiology |
| | BvH | 0.271 | 0.721 | | $4.042 \times 10^{-3}$ | 0.98219 | -4.3564 | | | | | |
| | Leak | 0.196 | 0.871 | $7.08 \times 10^{-4}$ | $1.72 \times 10^{-3}$ | 0.99734 | 4.2225 | | | | | |
| | Turgor | 0.2 | 0.864 | $4.07 \times 10^{-1}$ | $1.386 \times 10^{-3}$ | 0.99786 | 2.7128 | | | | | |
| 6,3 | BvH (Hyper) | 0.169 | 0.818 | | | | | Monkey | kidney fibroblast | 5 | 0.313<br>1.96 | Peckys and Mazur<br>2012.<br>Cryobiology |
| | BvH | 0.302 | 0.603 | | $5.922 \times 10^{-3}$ | 0.93913 | 19.098 | | | | | |
| | Leak | 0.156 | 0.844 | $2.23 \times 10^{-3}$ | $7.555 \times 10^{-4}$ | 0.99803 | ∞ | | | | | |
| | Turgor | 0.162 | 0.834 | 1.34 | $7.035 \times 10^{-4}$ | 0.99817 | ∞ | | | | | |
| 6,4 | BvH (Hyper) | 0.177 | 0.819 | | | | | Monkey | kidney fibroblast | 5 | 0.311<br>1.96 | Peckys et al.<br>2011.<br>PLoS ONE |
| | BvH | 0.29 | 0.643 | | $4.992 \times 10^{-3}$ | 0.95432 | 18.244 | | | | | |
| | Leak | 0.155 | 0.867 | $1.84 \times 10^{-3}$ | $4.181 \times 10^{-4}$ | 0.99898 | ∞ | | | | | |
| | Turgor | 0.161 | 0.857 | 1.08 | $3.085 \times 10^{-4}$ | 0.99925 | ∞ | | | | | |
| 7,1 | BvH (Hyper) | 0.355 | 0.645 | | | | | Rabbit | Corneal endothelial cell | 6 | 0.104<br>4. | Pegg et al.<br>1986.<br>Cell Biophysics |
| | BvH | 0.42 | 0.582 | | $1.151 \times 10^{-2}$ | 0.97718 | 8.6739 | | | | | |
| | Leak | 0.316 | 0.755 | $4.68 \times 10^{-4}$ | $3.63 \times 10^{-3}$ | 0.99691 | 32.475 | | | | | |
| | Turgor | 0.322 | 0.735 | $1.9 \times 10^{-1}$ | $2.699 \times 10^{-3}$ | 0.9977 | 30.697 | | | | | |
| 7,2 | BvH (Hyper) | 0.58 | 0.438 | | | | | Boar | Spermatozoa | 5 | 0.33<br>2. | Petrunkina et al.<br>2000.<br>Reproduction, Fertility and Development |
| | BvH | 0.632 | 0.348 | | $6.029 \times 10^{-4}$ | 0.9815 | 7.6751 | | | | | |
| | Leak | 0.578 | 0.44 | $9.77 \times 10^{-4}$ | $1.451 \times 10^{-4}$ | 0.99961 | ∞ | | | | | |
| | Turgor | 0.58 | 0.438 | 1.1 | $1.452 \times 10^{-4}$ | 0.99961 | ∞ | | | | | |
| 7,3 | BvH (Hyper) | 0.712 | 0.295 | | | | | Horse | Spermatozoa | 5 | 0.328<br>1.99 | Pommer et al.<br>2002.<br>Theriogenology |
| | BvH | 0.715 | 0.289 | | $1.055 \times 10^{-5}$ | 0.99952 | -12.554 | | | | | |
| | Leak | 0.71 | 0.296 | $6.48 \times 10^{-5}$ | $8.789 \times 10^{-6}$ | 0.99998 | ∞ | | | | | |
| | Turgor | 0.712 | 0.295 | $9.11 \times 10^{-2}$ | $8.789 \times 10^{-6}$ | 0.99998 | ∞ | | | | | |
| 7,4 | BvH (Hyper) | 0.755 | 0.253 | | | | | Monkey | Spermatozoa | 6 | 0.331<br>3.99 | Rutllant et al.<br>2003.<br>Journal of Andrology, |
| | BvH | 0.777 | 0.236 | | $1.337 \times 10^{-3}$ | 0.98171 | -4.2459 | | | | | |
| | Leak | 0.715 | 0.324 | $4.91 \times 10^{-4}$ | $3.623 \times 10^{-4}$ | 0.99945 | 18.648 | | | | | |
| | Turgor | 0.721 | 0.314 | $4.17 \times 10^{-1}$ | $2.298 \times 10^{-4}$ | 0.99965 | 15.916 | | | | | |
| 8,1 | BvH (Hyper) | 0.291 | 0.648 | | | | | Guinea-pigs | Myocyte | 5 | 0.501<br>2. | Sasaki et al.<br>1999.<br>Clinical and Experimental Pharmacology and Physiology |
| | BvH | 0.359 | 0.563 | | $1.262 \times 10^{-3}$ | 0.98211 | 11.367 | | | | | |
| | Leak | 0.253 | 0.709 | $8.8 \times 10^{-4}$ | $3.106 \times 10^{-4}$ | 0.99927 | ∞ | | | | | |
| | Turgor | 0.259 | 0.7 | $5.92 \times 10^{-1}$ | $2.164 \times 10^{-4}$ | 0.99949 | ∞ | | | | | |
| 8,2 | BvH (Hyper) | 0.38 | 0.632 | | | | | Bovine | Chondrocyte | 8 | 0.181<br>2.02 | Schugart<br>2005.<br>Thesis |
| | BvH | 0.388 | 0.613 | | $3.536 \times 10^{-4}$ | 0.99666 | -28.574 | | | | | |
| | Leak | 0.385 | 0.616 | $2.32 \times 10^{-7}$ | $3.536 \times 10^{-4}$ | 0.9994 | -18.782 | | | | | |
| | Turgor | 0.388 | 0.613 | $1.29 \times 10^{-4}$ | $3.537 \times 10^{-4}$ | 0.9994 | -18.781 | | | | | |
| 8,3 | BvH (Hyper) | 0.474 | 0.521 | | | | | Human | Fetal liver derived stem cell | 7 | 0.3<br>2.14 | Tarasov et al.<br>2004.<br>Cryobiology |
| | BvH | 0.536 | 0.425 | | $1.113 \times 10^{-3}$ | 0.98181 | -13.382 | | | | | |
| | Leak | 0.464 | 0.539 | $9.9 \times 10^{-4}$ | $6.756 \times 10^{-5}$ | 0.99989 | -18.435 | | | | | |
| | Turgor | 0.468 | 0.533 | $8.72 \times 10^{-1}$ | $4.316 \times 10^{-5}$ | 0.99993 | -21.571 | | | | | |
| 8,4 | BvH (Hyper) | 0.507 | 0.511 | | | | | Human | prostatic adenocarcinoma cells | 5 | 0.324<br>3.18 | Tatakamatsu et al.<br>2005.<br>Journal of Biomechanics |
| | BvH | 0.54 | 0.437 | | $2.108 \times 10^{-3}$ | 0.98651 | 13.934 | | | | | |
| | Leak | 0.538 | 0.439 | $1.83 \times 10^{-8}$ | $2.108 \times 10^{-3}$ | 0.9965 | ∞ | | | | | |
| | Turgor | 0.54 | 0.437 | $6.77 \times 10^{-6}$ | $2.108 \times 10^{-3}$ | 0.9965 | ∞ | | | | | |
| 9,1 | BvH (Hyper) | 0.161 | 0.851 | | | | | Human | HeLa | 12 | 0.289<br>2. | Tivey et al.<br>1985.<br>Membrane Biology |
| | BvH | 0.339 | 0.519 | | $7.578 \times 10^{-3}$ | 0.90148 | -15.348 | | | | | |
| | Leak | 0.158 | 0.853 | $7.08 \times 10^{-3}$ | $1.097 \times 10^{-4}$ | 0.99979 | -61.198 | | | | | |
| | Turgor | 0.163 | 0.845 | 4.87 | $1.108 \times 10^{-4}$ | 0.99979 | -61.074 | | | | | |
| 9,2 | BvH (Hyper) | 0.69 | 0.322 | | | | | Medaka | Oocyte MII | 5 | 0.158<br>1.99 | Valdez et al.<br>2005.<br>Cryobiology |
| | BvH | 0.745 | 0.206 | | $2.073 \times 10^{-3}$ | 0.87266 | 13.849 | | | | | |
| | Leak | 0.687 | 0.326 | $5.9 \times 10^{-3}$ | $6.65 \times 10^{-4}$ | 0.99824 | ∞ | | | | | |
| | Turgor | 0.689 | 0.323 | 7.78 | $6.606 \times 10^{-4}$ | 0.99825 | ∞ | | | | | |
| 9,3 | BvH (Hyper) | 0.474 | 0.532 | | | | | Medaka | Oocyte GV | 5 | 0.22<br>2.26 | Valdez et al.<br>2005.<br>Cryobiology |
| | BvH | 0.393 | 0.705 | | $5.797 \times 10^{-3}$ | 0.9738 | 18.991 | | | | | |
| | Leak | 0.389 | 0.708 | $3.09 \times 10^{-8}$ | $5.797 \times 10^{-3}$ | 0.99182 | ∞ | | | | | |
| | Turgor | 0.392 | 0.706 | $4.22 \times 10^{-3}$ | $5.796 \times 10^{-3}$ | 0.99182 | ∞ | | | | | |
| 9,4 | BvH (Hyper) | 0.101 | 0.911 | | | | | Bovine | Oocyte MII | 10 | 0.219<br>2.15 | Wang et al.<br>2010.<br>Cryobiology |
| | BvH | 0.261 | 0.579 | | $9.56 \times 10^{-3}$ | 0.92724 | -7.8917 | | | | | |
| | Leak | 0.102 | 0.902 | $3.63 \times 10^{-3}$ | $9.431 \times 10^{-4}$ | 0.99835 | -24.718 | | | | | |
| | Turgor | 0.108 | 0.891 | 2.57 | $9.501 \times 10^{-4}$ | 0.99834 | -24.643 | | | | | |
| 10,1 | BvH (Hyper) | 0.579 | 0.446 | | | | | Mouse | Spermatozoa | 5 | 0.233<br>3.86 | Willoughby et al.<br>1996.<br>Biology of Reproduction |
| | BvH | 0.613 | 0.393 | | $1.38 \times 10^{-3}$ | 0.99316 | 11.814 | | | | | |
| | Leak | 0.581 | 0.44 | $1.48 \times 10^{-4}$ | $1.025 \times 10^{-3}$ | 0.9985 | ∞ | | | | | |
| | Turgor | 0.587 | 0.43 | $8.19 \times 10^{-2}$ | $1.041 \times 10^{-3}$ | 0.99848 | ∞ | | | | | |
| 10,2 | BvH (Hyper) | 0.446 | 0.572 | | | | | Canine | Pancreatic islet cell | 5 | 0.249<br>4. | Woods et al.<br>1997.<br>Cryobiology |
| | BvH | 0.594 | 0.355 | | $8.372 \times 10^{-3}$ | 0.95392 | 20.829 | | | | | |
| | Leak | 0.435 | 0.594 | $9.62 \times 10^{-4}$ | $4.471 \times 10^{-4}$ | 0.99928 | ∞ | | | | | |
| | Turgor | 0.45 | 0.565 | $5.43 \times 10^{-1}$ | $4.004 \times 10^{-4}$ | 0.99935 | ∞ | | | | | |
| 10,3 | BvH (Hyper) | 0.394 | 0.606 | | | | | Canine | Pancreatic islet cell | 5 | 0.249<br>4. | Woods et al.<br>1997.<br>Cryobiology |
| | BvH | 0.553 | 0.376 | | $1.008 \times 10^{-2}$ | 0.95085 | 21.76 | | | | | |
| | Leak | 0.375 | 0.644 | $1.02 \times 10^{-3}$ | $1.784 \times 10^{-4}$ | 0.99971 | ∞ | | | | | |
| | Turgor | 0.39 | 0.614 | $5.47 \times 10^{-1}$ | $9.828 \times 10^{-6}$ | 0.99998 | ∞ | | | | | |
| 10,4 | BvH (Hyper) | 0.323 | 0.681 | | | | | Human | Pancreatic islet cell | 5 | 0.251<br>3.99 | Woods et al.<br>1997.<br>Cryobiology |
| | BvH | 0.535 | 0.373 | | $1.801 \times 10^{-2}$ | 0.9132 | 24.658 | | | | | |
| | Leak | 0.295 | 0.741 | $1.5 \times 10^{-3}$ | $5.489 \times 10^{-4}$ | 0.99909 | ∞ | | | | | |
| | Turgor | 0.31 | 0.71 | $8.16 \times 10^{-1}$ | $1.324 \times 10^{-4}$ | 0.99978 | ∞ | | | | | |
| 11,1 | BvH (Hyper) | 0.255 | 0.751 | | | | | Human | Pancreatic islet cell | 5 | 0.249<br>4. | Woods et al.<br>1997.<br>Cryobiology |
| | BvH | 0.474 | 0.421 | | $1.582 \times 10^{-2}$ | 0.9389 | 24.011 | | | | | |
| | Leak | 0.245 | 0.769 | $1.22 \times 10^{-3}$ | $5.043 \times 10^{-5}$ | 0.99992 | ∞ | | | | | |
| | Turgor | 0.263 | 0.733 | $5.91 \times 10^{-1}$ | $6.542 \times 10^{-5}$ | 0.9999 | ∞ | | | | | |
| 11,2 | BvH (Hyper) | 0.25 | 0.754 | | | | | Human | Dermal fibroblast cell | 10 | 0.216<br>2.15 | Yang et al.<br>2007.<br>Cell preservation technology |
| | BvH | 0.367 | 0.509 | | $6.771 \times 10^{-3}$ | 0.93332 | -11.341 | | | | | |
| | Leak | 0.254 | 0.739 | $2.57 \times 10^{-3}$ | $1.948 \times 10^{-3}$ | 0.99671 | -17.466 | | | | | |
| | Turgor | 0.26 | 0.727 | 2.02 | $1.919 \times 10^{-3}$ | 0.99676 | -17.615 | | | | | |
| 11,3 | BvH (Hyper) | 0.553 | 0.506 | | | | | Human | Erythrocyte | 13 | 0.0948<br>2.52 | Zhao et al.<br>2004.<br>Biophysical Chemistry |
| | BvH | 0.536 | 0.535 | | $4.071 \times 10^{-3}$ | 0.97031 | -25.82 | | | | | |
| | Leak | 0.534 | 0.537 | $2.54 \times 10^{-9}$ | $4.071 \times 10^{-3}$ | 0.99414 | -21.247 | | | | | |
| | Turgor | 0.536 | 0.535 | $1.64 \times 10^{-6}$ | $4.071 \times 10^{-3}$ | 0.99414 | -21.247 | | | | | |
| 11,4 | BvH (Hyper) | 0.174 | 0.84 | | | | | Hamster | fibroblast cell | 8 | 0.15<br>2.01 | Muldrew et al.<br>2009.<br>Cryobiology |
| | BvH | 0.188 | 0.804 | | $2.673 \times 10^{-4}$ | 0.99878 | -30.811 | | | | | |
| | Leak | 0.169 | 0.848 | $2.54 \times 10^{-4}$ | $5.706 \times 10^{-5}$ | 0.99989 | -33.375 | | | | | |
| | Turgor | 0.173 | 0.842 | $1.27 \times 10^{-1}$ | $5.415 \times 10^{-5}$ | 0.9999 | -33.794 | | | | | |
