## Supplemental Material 2 (Group 2 Data) for "Meta-analysis and experimental re-evaluation of the Boyle van ‘t Hoff relation with osmoregulation modelled by linear elastic principles and ion-osmolyte leakage"

| DW score | DW crit | Animal Type | Cell Type | # data points | Min & Max Range | Journal Article |
|---|---|---|---|---|---|---|
| 1.96182 | 0.61 | Rat | Synaptosomes | 6 | 0.657<br>1. | Babila et al.<br>1990.<br>Journal of Neurochemistry |
| 3.29109 | 0.61 | Human | Chondrocyte | 6 | 0.542<br>1. | Bush and Hall<br>2005.<br>Journal of cellular physiology |
| 1.67061 | 0.61 | Boar | Spermatozoa | 6 | 0.194<br>1. | Du et al.<br>1994.<br>Theriogenology |
| 2.98297 | 0.61 | Human | Chondrocyte | 6 | 0.537<br>0.955 | Guilak<br>2000.<br>Bioreheology |
| 0.500881 | 0.879 | Human | Erythrocyte | 10 | 0.0979<br>0.994 | Heubusch et al.<br>1985.<br>Journal of cellular physiology |
| 0.657551 | 0.879 | Human | Erythrocyte | 10 | 0.184<br>0.996 | Heubusch et al.<br>1985.<br>Journal of cellular physiology |
| 1.41029 | 0.61 | Human | CD34+ cell | 6 | 0.161<br>1. | Hunt et al.<br>2003.<br>Cryobiology |
| 2.14794 | 1.077 | Human | Lymphocyte | 15 | 0.103<br>0.986 | Law et al.<br>1983.<br>Cryobiology |
| 1.53769 | 0.824 | Human | Lymphocyte | 9 | 0.381<br>1.03 | McGann et al.<br>1988.<br>Cytometry |
| 1.8599 | 0.763 | Human | Granulocyte | 8 | 0.399<br>0.999 | McGann et al.<br>1988.<br>Cytometry |
| 2.89778 | 0.763 | Hamster | Fibroblast | 8 | 0.398<br>0.997 | McGann et al.<br>1988.<br>Cytometry |
| 2.00076 | 0.7 | Sea Urchin | Oocyte | 7 | 0.448<br>1.02 | Mela<br>1967.<br>Biophysical Journal |
| 2.00502 | 0.763 | Mouse | Ova | 8 | 0.0846<br>0.983 | Oda et al.<br>1992.<br>Journal of reproduction and fertility |
| 3.40792 | 0.61 | Bovine | Chondrocyte | 6 | 0.181<br>0.999 | Schugart<br>2005.<br>Thesis |
| 2.42631 | 0.7 | Human | Jurkat cell | 7 | 0.432<br>0.878 | Simmons<br>1984.<br>Journal of Experimental Physiology |
| 2.26016 | 0.763 | Human | HeLa | 8 | 0.289<br>1. | Tivey et al.<br>1985.<br>Membrane Biology |
| 2.07136 | 0.7 | Bovine | Oocyte GV | 7 | 0.316<br>1.04 | Wang et al.<br>2010.<br>Cryobiology |
| 1.84992 | 0.61 | Bovine | Oocyte MII | 6 | 0.219<br>1. | Wang et al.<br>2010.<br>Cryobiology |
| 2.97889 | 0.61 | Human | Dermal fibroblast cell | 6 | 0.216<br>0.997 | Yang et al.<br>2007.<br>Cell preservation technology |
| 2.59387 | 0.61 | Human | Osteoblast | 6 | 0.123<br>0.998 | Yang et al.<br>2011.<br>Journal of Chemical Engineering of Chinese Universities |
| 1.49751 | 0.879 | Human | Erythrocyte | 10 | 0.0948<br>1. | Zhao et al.<br>2004.<br>Biophysical Chemistry |
| 2.01852 | 0.61 | Hamster | fibroblast cell | 6 | 0.15<br>1. | Muldrew et al.<br>2009.<br>Cryobiology |
