## Supplemental Material 1 (Group 1 Data) for "Meta-analysis and experimental re-evaluation of the Boyle van ‘t Hoff relation with osmoregulation modelled by linear elastic principles and ion-osmolyte leakage"

| DW score | DW crit | Animal Type | Cell Type | # data points | Min & Max Range | Journal Article |
|---|---|---|---|---|---|---|
| 1.28738 | 0.61 | Golden Hamster | Pancreatic islet cell | 6 | 0.15<br>2.01 | Benson et al. 1993.<br>Cell Transplantation |
| 3.17941 | 0.7 | Bovine | Spermatozoa | 7 | 0.502<br>3. | Basse et al. 2012.<br>Theriogenology |
| 0.736696 | 0.927 | Human | Chondrocyte | 11 | 0.542<br>9.53 | Bush and Hall 2005.<br>Journal of cellular physiology |
| 1.11159 | 0.763 | Lobster | axon | 8 | 0.32<br>4. | Freeman et al. 1966.<br>The Journal of general physiology |
| 1.17065 | 0.763 | Human | hematopoietic progenitor cells | 8 | 0.215<br>1.99 | Gao et al. 1998.<br>Cryobiology |
| 2.68093 | 0.824 | Human | Chondrocyte | 9 | 0.537<br>3. | Guilak 2000.<br>Bioreheology |
| 1.04258 | 0.879 | Human | CD34+ cell | 10 | 0.161<br>1.95 | Hunt et al. 2003.<br>Cryobiology |
| 0.803726 | 0.763 | Dog | notochordal cell | 8 | 0.335<br>7.81 | Hunter et al. 2007.<br>Mol Cell Biomech. |
| 0.867032 | 0.879 | Crab | Muscle Fiber | 10 | 0.33<br>2.07 | Lang and Gainer 1969.<br>The Journal of General Physiology |
| 2.35341 | 1.133 | Human | Lymphocyte | 17 | 0.103<br>2.03 | Law et al. 1983.<br>Cryobiology |
| 1.15163 | 0.61 | Hamster | Pancreatic islet cell | 6 | 0.201<br>3.35 | Lui et al. 1995.<br>Cryobiology |
| 1.42482 | 0.7 | Mouse | Blastocyte (with zona) | 7 | 0.174<br>4.93 | Mazur and Schneider 1985.<br>Cell Biophysics |
| 1.66377 | 0.879 | Bovine | Morulae | 10 | 0.118<br>2.86 | Mazur and Schneider 1985.<br>Cell Biophysics |
| 1.24258 | 1.101 | Mouse | Ova | 13 | 0.0846<br>5.6 | Oda et al. 1992.<br>Journal of reproduction and fertility |
| 1.71272 | 0.7 | Horse | Spermatozoa | 7 | 0.498<br>2. | Oldenhof et al. 2011.<br>Theriogenology |
| 0.47183 | 0.824 | Hamster | Oocyte (with zona) | 9 | 0.0471<br>5.99 | Oungoulian et al. 2011.<br>Proceedings of the ASME 2011 Summer Bioengineering Conference |
| 2.1501 | 0.7 | Mouse | cortex neuron | 7 | 0.23<br>2.31 | Paynter et al. 2009.<br>Cryobiology |
| 2.2455 | 0.7 | Mouse | cortex neuron | 7 | 0.232<br>2.32 | Paynter et al. 2009.<br>Cryobiology |
| 2.3588 | 0.7 | Mouse | ventral mesencephalon nerual cell | 7 | 0.232<br>2.32 | Paynter et al. 2009.<br>Cryobiology |
| 2.10431 | 0.61 | Rabbit | Corneal endothelial cell | 6 | 0.104<br>4. | Pegg et al. 1986.<br>Cell Biophysics |
| 2.51105 | 0.61 | Monkey | Spermatozoa | 6 | 0.331<br>3.99 | Rutllant et al. 2003.<br>Journal of Andrology, |
| 2.91594 | 0.763 | Bovine | Chondrocyte | 8 | 0.181<br>2.02 | Schugart 2005.<br>Thesis |
| 0.964349 | 0.7 | Human | Fetal liver derived stem cell | 7 | 0.3<br>2.14 | Tarasov et al. 2004.<br>Cryobiology |
| 0.481465 | 0.971 | Human | HeLa | 12 | 0.289<br>2. | Tivey et al. 1985.<br>Membrane Biology |
| 0.805827 | 0.879 | Bovine | Oocyte MII | 10 | 0.219<br>2.15 | Wang et al. 2010.<br>Cryobiology |
| 1.45293 | 0.879 | Human | Dermal fibroblast cell | 10 | 0.216<br>2.15 | Yang et al. 2007.<br>Cell preservation technology |
| 1.34672 | 1.101 | Human | Erythrocyte | 13 | 0.0948<br>2.52 | Zhao et al. 2004.<br>Biophysical Chemistry |
| 1.74968 | 0.763 | Hamster | fibroblast cell | 8 | 0.15<br>2.01 | Muldrew et al. 2009.<br>Cryobiology |
